# The N-end Rule Pathway and Ubr1 enforce protein compartmentalization via N-terminally-encoded cellular location signals

**DOI:** 10.1101/392373

**Authors:** Anthony Tran

## Abstract

The Arg/N-end rule pathway, a mechanism of protein degradation conserved from yeast to humans, is involved in cellular protein quality control, but its role has only been vaguely understood. Through systematic examination of single residue mutants of model misfolded substrates, and global analyses of yeast proteins, we discovered that Ubr1, an E3 ligase of the Arg/N-end rule, degrades organellar proteins that fail to reach their intended subcellular compartments. We determined that recognition by Ubr1 is dependent on location signals that are naturally embedded into the 2nd amino acid residue of the majority of proteins. The N-end rule pathway is thus likely to have been critical to the evolution of endosymbiotic relationships which paved the way for advanced eukaryotic cellular life.

**Significance Statement:** This work elucidates a novel role for the N-end Rule Pathway, a protein degradation pathway highly conserved from yeast to humans. We demonstrate that the N-end rule pathway enforces the cellular compartmentalization of ER and mitochondrial proteins by degrading them when they fail to successfully translocate into their intended destinations and thus become mislocalized to the cytosol. This mechanism prevents the accumulation of toxic foreign proteins within the cytosol. Recognition of the displaced proteins is dependent on cellular location signals programmed into the 2^nd^ residue of the target proteins, as well as the tendency for the proteins to misfold in a foreign environment. These findings have significant relevance to research on the mechanisms causing human diseases involving protein misfolding.

## Main Text

Protein quality control (PQC) is an essential protein quality surveillance and degradation system through which cells ensure the integrity of the proteome and maintain cellular homeostasis *(1)*. Uncontrolled aggregation of misfolded proteins leads to the formation of insoluble protein deposits that are detrimental to the cell and are widely associated with numerous human diseases such as Alzheimer’s, Parkinson’s, and Huntington’s disease *(2–4)*. Cytosolic quality control (CytoQC) pathways, a subset of PQC, specifically mediates the clearance of cytosolically-localized aberrant proteins. In yeast, several E3s have been implicated in CytoQC pathways with varying client substrate scopes and enzymatic activities that are in several cases also dependent on specific physiological insults, such as heat or chemical stresses *(5–9)*. The San1 and Ubr1 E3 ligases have partially overlapping specificities in the degradation of several cytosolic model misfolded substrates *(7,8)*. San1p targets subsrates with exposed hydrophobicity consisting of contiguous sequences of at least five or more hydrophobic residues, and was first identified as an E3 ligase that mediates quality control of nuclear proteins *(10–11)*. Ubr1p is best known for its role as an N-recognin in the N-end rule pathway, which relates the half-life of a protein to its N-terminal residue *(12)*. Ubr1 possesses two domains, the UBR box and N-domain, which recognize and bind to proteins with compatible N-terminal residues, leading to their ubiquitin-dependent degradation. In eukaryotes, three branches of the N-end rule pathway exist: Arg/N-end rule, Ac/N-end rule, and the more recently characterized Pro/N-end rule. Ubr1p mediates the Arg/N-end branch, while two other E3 ligases, Doa10p and Gid4p, are N-recognins of the latter two branches, respectively. Together, these distinct branches have been been demonstrated to partake in regulating a wide range of biological processes, such as oxygen and nitric oxide sensing, chromosome segregation and DNA replication, autophagy, cell migration and neurogenesis, and many others *(13)*.

While there had been earlier conflicting reports on the role of the N-end rule pathway in Ubr1-mediated CytoQC*(8,14)*, it was later demonstrated that ΔssC^22-519^Leu2_myc_, a model misfolded cytosolic substrate, was recognized by Ubr1 through its N-terminal Met-Ile, a Met-Φ N-degron, defined as an N-terminal Met followed by a Leu, Phe, Tyr, Trp, or Ile residue at the P2 position *(15,16)*. The latter finding definitively linked Ubr1-mediated CytoQC with N-degrons of the Arg/N-end rule pathway for at least a subset of substrates. A role in protein quality control has also been demonstrated for the Ac/N-end branch of the pathway in the case of orphaned protein subunits that fail to merge into their native complexes due to stoichiometric imbalances *(17)*.

Despite the recent confirmations that recognition of some misfolded substrates in the cytosol by Ubr1 is N-terminal sequence dependent, the full scope and underlying physiological role of the Arg/N-end rule pathway in the context of CytoQC is not clearly understood. The P2 position amino acid residue is a primary determinant of whether leading methionine excision of a nascent protein occurs *(18,19)*, and thus, is also the deciding factor in the final native N-terminal sequence of a protein prior to downstream processing steps such as acetylation or signal sequence cleavage. Therefore, absent other N-terminal modifications such as acetylation that may block recognition of the N-terminus, the P2 residue of a protein is the primary determinant of whether the native unmodified N-terminal sequence of a nascent protein is compatible with recognition by the Arg/N-end rule pathway after translation in the cytosol. Here we successfully characterize the set of P2 amino acid residues which produce an expanded set of Ubr1 compatible N-terminal N-degrons specific to CytoQC by conducting a systematic biochemical analyses of P2-residue variants of a model misfolded CytoQC substrate, Ste6*C *(20)*. This expanded set of N-degrons differs from the original set of destabilizing Arg/N-end rule residues, which include primary residues, single basic or bulky hydrophobic residues exposed at the N-terminus, and secondary and tertiary residues, to which a primary destabilizing residue is appended as a result of one or two step modification processes *(12)*. Instead, the expanded N-degron set we characterize here closely overlaps with the more recently discovered set of Met-Φ N-degrons, which involve a leading methionine residue followed by a compatible P2 residue *(15)*. While the originally delineated Arg/N-end rule destabilizing residues generally do not exist at the N-termini of proteins in their native cytosolic or pre-translocated forms because they prevent leading methionine cleavage due to their size *(18,19)*, Met-Φ N-degrons are naturally abundant, as all proteins are first translated with a leading methionine. Our approach allowed us to identify, in vivo, additional N-degrons relevant to misfolded proteins that were not previously detected using peptide arrays on membrane support (SPOT), which only utilizes short peptides for in vitro binding assays *(15)*. Next, we performed detailed global bioinformatic analysis of P2 amino acid usage frequency in the *S. cerevisiae* proteome, taking into account the expanded P2-dependent N-degron set, and found that Ubr1’s specificity is designed to preferentially degrade secretory and mitochondrial proteins that fail to translocate and thus become mislocalized to the cytosol. Our precise in vivo analyses of turnover rates of several endogenous translocation-inhibited secretory and mitochondrial proteins in the presence and absence of Ubr1 confirm this model of CytoQC activity. Finally, an in-depth analysis of a representative set of signal-sequence bearing proteins revealed that ∼93% of soluble ER proteins are encoded with Ubr1-QC-compatible P2-residues, versus only ∼26% of natively localized cytosolic proteins, indicating that P2-residues encode signals of cellular location that also facilitate the degradation of displaced proteins in the cytosol. Taken together, the findings demonstrate that the N-terminal P2-encoding of cellular location, and consequently, the N-end rule pathway, both serve essential roles in enforcing the fidelity of protein compartmentalization in eukaryotic cells.

## Results

### P2-residue identity enables or inhibits degradation of misfolded substrates by Ubr1

Ste6*C is a truncated form of Ste6p, a plasma membrane ATP-binding cassette transporter, containing only the cytosolic tail of the protein, and harbors a deletion of residues 1249-1290. Δ2GFP is a derivative of wild-type GFP wherein residues 25-36 are deleted. The deletions in each substrate cause misfolding, resulting in their degradation by CytoQC pathways*(7,20)*. Efficient degradation of each substrate requires both Ubr1 and San1 E3 ligases with differing dependencies: the degradation of Ste6*C is biased towards the Ubr1 pathway, whereas the degradation of Δ2GFP is biased towards the San1 pathway **(Figures 1A, 1B)**. Based on the rules of N-terminal methionine excision, Ste6*C is predicted to possess a Met-Ile sequence at the N-terminus, while Δ2GFP would harbor a -Ser upon methionine excision *(18,19)*. N-terminal Met-Ile, an N-degron of the Arg/N-end rule pathway, should help direct Ste6*C towards the Ubr1 pathway, while Δ2GFP, with an N-terminal -Ser, is shielded from Ubr1 N-degron recognition *(12,15)*. Since the N-terminal sequence of a misfolded protein impacts its recognition by the Ubr1 pathway, we surmised that by altering the N-terminal sequences of Ste6*C and Δ2GFP, it would be possible to increase or decrease the amount of degradation which proceeds via Ubr1. We thus generated P2 mutants of Ste6*C and Δ2GFP that matched P2-residue of the other to assess the impact on Ubr1-mediated degradation. We modified the N-terminus of Ste6*C to mimic that of Δ2GFP by substituting the P2 position Ile of Ste6*C (Ste6*C-I2I) with a Ser residue, generating Ste6*C-I2S, which carries an N-terminal -Ser. This was confirmed through Edman’s degradation sequencing of immunoprecipitated protein **(Figure S1A).** In contrast to Ste6*C-I2I, the presence of Ubr1p did not increase degradation rates of Ste6*C-I2S in either Δ*san1* or + SAN1 backgrounds (Δ*san1*Δ*ubr1* vs Δ*san1* + UBR1; +SAN1Δ*ubr1* vs + SAN1 + UBR1) (**Figure 1C**). Thus, by substituting the P2 Ile of Ste6*C with Ser, Ubr1-mediated degradation of Ste6*C was effectively inhibited. On the other hand, degradation of Ste6*C-I2S in the presence of only San1p remained similar to Ste6*C-I2I **(Figures 1A, 1C**; +SAN1Δ*ubr1*).

**Fig. 1.**
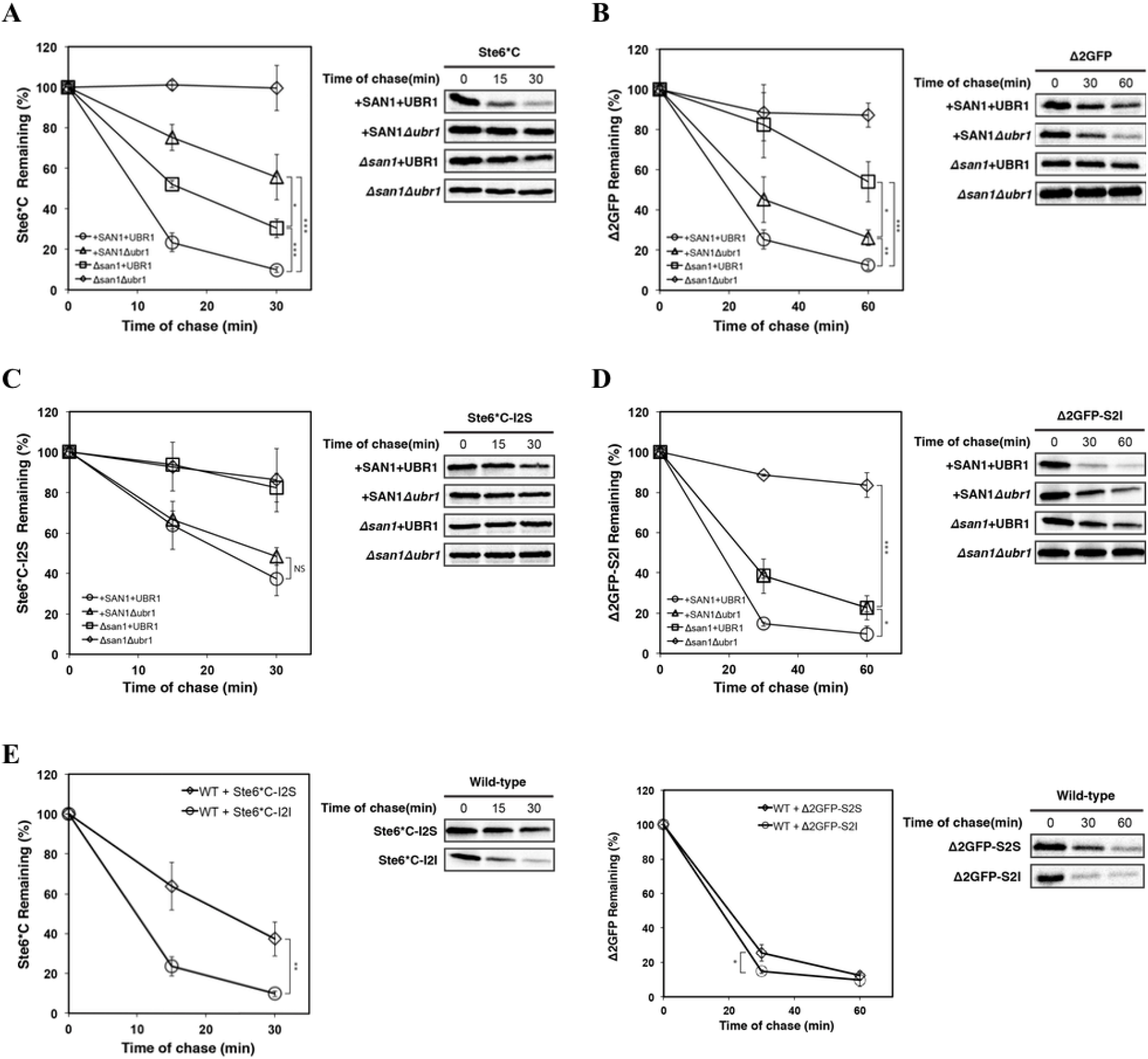
Efficient degradation of Ste6*C and Δ2GFP by the Ubr1 quality control pathway is dependent on P2 residue identity. (A) and (B) Turnover of Ste6*C and Δ2GFP in + SAN1 + UBR1 (wild-type), Δsan1 + UBR1, +SAN1Δubr1, and Δsan1Δubr1 cells by metabolic pulse-chase analysis (C) and (D) Turnover of Ste6*C-I2S and Δ2GFP-S2I in + SAN1 + UBR1(wild-type), Δsan1 + UBR1, +SAN1Δubr1, and Δsan1Δubr1 cells was examined by metabolic pulse-chase analysis. (E) Comparison of turnover of P2-serine substrates (Ste6*C-I2S, Δ2GFP-S2S) with P2-isoleucine substrates (Ste6*C-I2I, Δ2GFP-S2I) in wild-type cells by metabolic pulse-chase analysis (A-E) Metabolic pulse-chase: cells were grown to log phase and shifted to 30°C for 30 minutes followed by pulse-labeling with [35S]methionine/cysteine (5 min for Ste*6C and 10 min for Δ2GFP) and chased at the times indicated. Proteins were immunoprecipitated using anti-HA antibody and resolved by SDS- PAGE, then visualized by phosphoimager analysis. Error bars, mean +/- SD of three independent experiments (N = 3, biological replicates). Student’s t-test: *, p < 0.05; **, p < 0.01; ***, p < 0.005; Not Significant (NS), p > 0.5.

The inverse experiment was performed with Δ2GFP (Δ2GFP-S2S) by substituting the P2 Ser with Ile, generating Δ2GFP-S2I. Degradation of Δ2GFP-S2I was far more efficient in Δ*san1* + UBR1 cells than Δ*san1*Δ*ubr1* cells, in contrast to Δ2GFP, for which the presence of Ubr1p (Δ*san1* + UBR1) conferred only a minimal increase of degradation efficiency (**Figures 1B and 1D**). As the case with Ste6*C-I2S, degradation efficiency of Δ2GFP via the San1-mediated pathway remained relatively unaffected by the S2I substitution (**Figures 1B and 1D**; +SAN1Δ*ubr1*). For both substrates, possessing a P2 Ile as opposed to a P2 Ser increased the overall degradation rate of the substrates in wild-type cells, suggesting that the enhancement of Ubr1-mediated degradation conferred by a P2 Ile occurs under normal physiological conditions (**Figure 1E**).

We next addressed whether the initiating methionine of Ste6*C-I2I was being cleaved through an unidentified mechanism specific to the processing of misfolded proteins, possibly presenting an exception to the known rules of N-terminal methionine excision. In the Arg/N-end rule pathway, isoleucine is a Type-2 destabilizing residue. Cleavage of the N-terminal methionine of Ste6*C-I2I would expose the isoleucine residue at the P2 position, triggering its recognition by Ubr1 and subsequent degradation via the Arg/N-end rule pathway. To determine if this was the case, we performed N-terminal sequencing on Ste6*C-I2I isolated from Δ*san1*Δ*ubr1* cells. This analysis confirmed that the leading methionine is retained by Ste6*C-I2I, as predicted **(Figure S1B)**. Therefore, recognition of Ste6*C and Δ2GFP-S2I by Ubr1p is through an N-terminal Met-Ile as opposed to an N-terminal Ile, both of which are Arg/N-end rule pathway N-degrons.

We observed that nine amino acids at the P2 position resulted in significant inhibition of Ubr1-mediated degradation of Ste6*C in Δ*san1* + UBR1 cells: Ala, Asn, Ser, Cys, Glu, Pro, Asp, Thr, and Met, which we refer to as Ubr1-QC-*incompatible* P2 amino acids. His, Trp, Phe, Tyr, Ile, Val, Gln, Gly, Lys, Leu, and Arg, allowed rapid degradation by Ubr1, which we refer to as Ubr1-QC-*compatible* P2 amino acids (**Figure 2A, S2A-S2C**). The same effect was not seen in + SAN1Δ*ubr1* cells, for which no statistically significant decrease of turnover rate relative to the original Ste6*C substrate was observed (**Figure 2A, S2D-S2F**). The Ubr1-dependent degradation of the 9 most rapidly degraded Ste6*C P2 mutants was confirmed in Δ*san1*Δ*ubr1* cells (**Figure S3**). Ste6*C P2 mutants inhibited for Ubr1 degradation was not a result of Ubr1 blockage by N-terminal acetylation since Ste6*C is encoded with a P3 Pro residue, which is known to prevent N-terminal acetylation *(21)*. Leading methionine retention is not a prerequisite for compatibility with the Ubr1-QC degradation pathway, as Ste6*C-I2 V was demonstrated to be degraded efficiently by Ubr1 and is predicted to undergo methionine cleavage, also confirmed via sequencing **(Figure S1C)**.

**Fig. 2.**
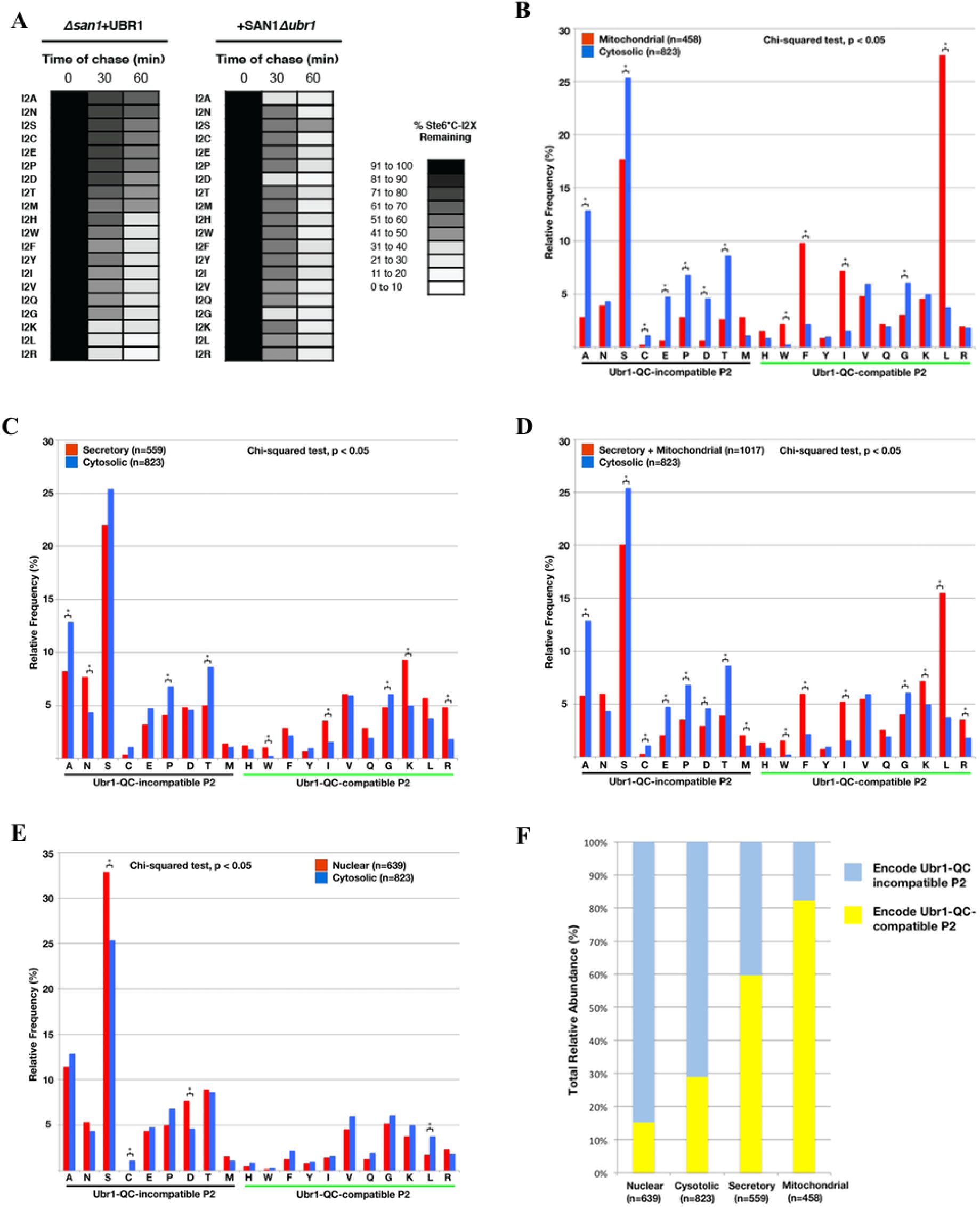
The P2-residue identity of a CytoQC substrate specifically effects degradation efficiency of the Ubr1 quality control pathway. Proteins localized to secretory pathway components and mitochondria exhibit bias for encoding UBR1-QC-compatible P2 residues when compared to cytosolically-localized proteins. (A) Turnover rates of nineteen Ste6*C P2 amino acid residue mutants (Ste6*C-I2X, where X = A[alanine], N[asparagine], S[serine], C[cysteine], E[glutamic acid], P[proline], D[aspartic acid], T[threonine], M[methionine], H[histidine], W[tryptophan], F[phenylalanine], Y[tyrosine], V[valine], Q[glutamine], G[glycine], K[lysine], L[leucine], R[arginine]) were compared to the turnover rate of the original Ste6*C substrate (Ste6*C-I2I) in Δsan1 + UBR1 and + SAN1Δubr1 cells by pulse chase analysis: cells were grown to log phase and shifted to 30°C for 30 minutes followed by pulse-labeling with [35S]methionine/cysteine for 5 minutes and chased at the times indicated. Proteins were immunoprecipitated using anti-HA antibody and resolved by SDS- PAGE, then visualized by phosphoimager analysis. Heat-map reflects results presented in Figures S2A, S2B, S2C, S2D, S2E, and S2F, and is based on the mean of three independent experiments (N = 3, biological experiments) unless otherwise indicated. (B) Relative frequency of P2 amino acid usage of proteins localized to the secretory pathway components (N = 559, proteins) vs cytosol (N = 823, proteins). Frequency distribution differed significantly between the two sets (p < 1 × 10^−6^, *x* ^2^ = 64.33, *df* = 19; Table S9). Relative usage frequency of amino acids Ala (A), Asn (N), Pro (P), Thr (T), Trp (W), Ile (I), Lys (K), and Arg (R) differed significantly between the two sets (pair-wise Chi-square analysis; p < 0.01 for A, N, K, R; p < 0.05 for P, T, W, I). (C) Relative frequency of P2 amino acid usage in proteins localized to mitochondria (N = 458, proteins) vs cytosol (N = 823, proteins). Frequency distribution differed significantly between the two sets (p < 1 × 10^−55^, *x* ^2^ = 317.85, *df* = 19; Table S9). Relative usage frequency of amino acids Ala (A), Ser(S), Cys (C), Glu (E), Pro (P), Asp (D), Thr (T), Trp (W), Phe (F), Ile (I), Gly (G), and Leu (L) differed significantly between the two sets (pair-wise Chi-square analysis; p < 0.0001 for A, S, E, P, D, T, F, I, L; p < 0.05 for C, W, G) (D) Relative frequency of P2 amino acid usage in proteins localized to either mitochondria or secretory pathway (N = 1017, proteins) vs cytosol (N = 823, proteins). Frequency distribution differed significantly between the two sets (p < 1 × 10^−30^, *x* ^2^ = 196.34, *df* = 19; Table S10). Relative usage frequency of amino acids Ala (A), Ser (S), Cys (C), Glu (E), Pro (P), Thr (T), Trp (W), Phe (F), Ile (I), Gly (G), Leu (L), Arg (R) differed significantly between the two sets (pair-wise Chi-square analysis; p < 0.0001 for A, T, F, I, L, R; p < 0.01 for S, E, P, W; p < 0.05 for C, G). (E) Relative frequency of P2 amino acid usage in proteins localized to nucleus (N = 639, proteins) vs cytosol (N = 823, proteins). Frequency distribution differed less significantly between the two sets (p > 0.005, *x*^2^ = 36.33, *df =* 19; Table S10). Relative usage frequency of amino acids Ser (S), Cys (C), Asp (D), Leu (L) differed significantly between the two sets (pair-wise Chi-square analysis; p < 0.001 for S; p < 0.01 for C, D, L). (F) Estimated total relative abundance of proteins that are encoded with Ubr1-QC-compatible P2 residues by localization category. Abundance levels based on Huh et al., 2003 *(22)*.

Interestingly, P2 His, Lys, Arg, and Gln, all of which cause retention of initiating methionines based on the conventional rules of methionine excision, are not predicted to produce N-termini with known N-degrons of the N-end rule pathway *(12,15,18,19)*. However, it is important to note that unconventional N-terminal methionine excision has been demonstrated to occur for some reporter substrates and endogenous proteins with P2 Asn, Gln, and His residues, for which Ubr1-mediated turnover was discovered to be reliant on the atypical removal of the initiating methionine residues, and resultant exposure, Nt-amidation and Nt-arginylation of the P2 residues in the case of P2 Asn and Gln, or simply the exposure of the P2 residue in the case of P2 His. *(49,50,51)*. It is therefore also a possibility that the Ste6*C mutants in this study possessing a P2 His, Lys, Arg, or Gln, undergo non-canonical methionine cleavage reactions, resulting in N-terminally exposed His, Lys or Arg, Type 2 primary destabilizing residues of the N-end rule pathway, or N-terminally exposed Gln, a tertiary destabilizing residue that can be converted into a primary destabilizing Arg residue through N-terminal amidation followed by N-terminal arginylation *(12)*. Unconventional initiating methionine removal mechanisms in the case of P2 Lys, and Arg residues have yet to be demonstrated. Thus, our experiments, combined with current knowledge of N-terminal methionine excision pathways, demonstrate that P2 His, Lys, Arg, and Gln residues may either induce non-conventional N-terminal methionine excision that then enable their processing and targeting through the canonical N-end rule, or alternatively, generate novel N-terminal Arg/N-end rule N-degrons that are only recognizable by Ubr1 when present on substrates that are misfolded. Furthur analysis of these individual Ste6*C variants would be required to confirm which specific processing mechanisms, if any, lead to their recognition and degradation by Ubr1.

P2 Gly and Val residues are predicted to permit leading methionine removal and become exposed at the N-terminus. However, N-terminal Gly and Val have thus far not been characterized as N-degrons. Interestingly, Met-Gly and Met-Val N-terminal sequences have also not been previously demonstrated to function as N-degrons. Therefore, even in the unexpected events where the leading methionines are somehow retained for these variants, our experiments with Ste6*C suggest that Gly and Val encoded at the P2 position result in novel N-degrons that are compatible with Ubr1-mediated degradation.

That the Ubr1-compatibility of P2 His, Lys, Arg, Gln, with the initiating methionine retained, or P2 Gly and Val, with the initiating methionine removed, was not detected in assays involving short peptides in previously published studies indicates that additional allosteric interaction with Ubr1 may be taking place in the case of misfolded substrates that increases the range of Ubr1’s specificity for N-degron sequences. The exact reason for this extended specificity could be an area of further study. Once again, an alternative explanation for the compatibility of P2 His, Lys, Arg, or Gln with Ubr1 in our experiments is that non-conventional N-terminal methionine excision, and consequent exposure of the P2 residues, occurs under physiological conditions, but does not occur in *in vitro* assays utilizing short peptides *(15).* It is also worth noting that in a high-throughput analysis of N-degrons utilizing mCherry-sfGFP as a reporter substrate, while reporters with Met-Φ degrons were on average less stable than other variants, the observed destabilization was independent of Ubr1. In that study, reporters with P2 His, Lys, Arg, Gln, Gly, and Val were either moderately or highly stable. The lack of correlation between the results obtained with the mCherry-sfGFP reporter substrate used in their anlaysis, and those achieved using our misfolded substrates, Ste6*C and Δ2GFP, demonstrates again that conformational aberrancy is a key determinant in N-degron compatibility with Ubr1 *(49)*.

Interestingly, the mutation of P2 Leu to Lys in the case of ΔssC^22-519^Leu2_myc_ inhibited its Ubr1-mediated degradation *(15)*, in contrast to Ste6*C, for which both P2 Leu and Lys were compatible with Ubr1 degradation (**Figure 2A**). While a concomitant change in acetylation state was not ruled out for ΔssC^22-519^Leu2_myc_ as a result of the L2K mutation, which would block N-terminal recognition by Ubr1, the difference in outcomes suggest that there are indeed context and substrate specific influences on the Arg/N-end rule pathway’s activity in CytoQC that need to be further dissected. Thus far, the expanded set of N-degrons discovered using Ste6*C is expected to apply generally to misfolded proteins in the cytosol, but may extend to other targets and physiological states not explored here.

### The majority of secretory and mitochondrial proteins encode Ubr1-QC-compatible P2-residues

To determine if there is a bias within specific cellular compartments for proteins to possess P2 residues that are Ubr1-QC-compatible, we utilized data from a genome-wide GFP-fusion based localization study of protein localization spanning 4156 proteins *(22)* **(Table S2-S4)**. P2 amino acid usage frequency was analyzed for proteins with exclusive nuclear, cytosolic, secretory pathway, or mitochondrial localization (**Tables S5-S8; refer to Materials and Methods**). Frequency distribution across all residues was significantly different for secretory (n = 559, p < 1 × 10^−6^, *x* ^2^ = 64.33, *df* = 19) and mitochondrial proteins (n = 458, p < 1 × 10^−55^, *x* ^2^ = 317.85, *df* = 19) when compared against cytosolic proteins (n = 823) **(Figures 2B-D; Tables S9 and S10)**. Pair-wise chi-square analysis of individual amino acid usage frequencies demonstrated that biases were seen for the mitochondrial and secretory proteins to be encoded with P2 residues that are Ubr1-QC-compatible, with 6 Ubr1-QC-compatible P2 amino acids (Trp, Phe, Ile, Lys, Leu, and Arg) having statistically significant higher percentage usage frequencies in at least one or both of the mitochondrial and secretory pathway protein sets when compared with cytosolic proteins. An equally important finding is that we observed a significant usage bias *against* 7 amino acids that do *not* enable efficient Ubr1 degradation (Ubr1-QC-*incompatible* P2 residues Ala, Ser, Cys, Glu, Pro, Asp, Thr) in either one or both mitochondrial and secretory pathway protein sets when compared to cytosolic proteins. In contrast, frequency distribution between nuclear and cytosolic proteins had a much less significant difference (nuclear, n = 639; cytosolic n = 823; p > 0.001, *x*^2^ = 36.33, *df* = 19), and only four significant differences in relative frequencies of individual amino acids, and no general bias for or against Ubr1-QC P2 compatible residues (**Figure 2E; Table S10**).

We determined the total relative abundance of proteins within each localization category based on abundance level data gathered from the global yeast protein GFP-fusion study conducted by Huh and colleagues *(22)* (refer to Materials and Methods). The majority of nuclear and cytosolically localized protein is encoded with Ubr1-QC-*incompatible* P2 amino acids (84.7% and 70.9%, respectively), while the majority of secretory and mitochondrial protein is associated with Ubr1-QC-*compatible* P2 amino acids (59.7% and 82.3%, respectively) (**Figure 2F; Table S17-S27**).

The much higher prevalence of Ubr1-QC-compatible proteins in the latter two categories indicated that Ubr1p may be optimized for the degradation of secretory and mitochondrial protein. That they do not natively reside in the cytosol suggested a model in which proteins in these pathways are subject to Ubr1-mediated CytoQC when they fail to translocate. Compartment-specific chaperones and enzymes are important for the native folding processes of secretory and mitochondrial proteins *(23–25)*. Spontaneous attempts at folding by mitochondrial proteins in the cytosol have been proven to occur, thereby preventing import*(26).* Various cytosolic chaperones are designed to maintain pre-translocated proteins in partially-folded, import-competent states, while the folding efficiency of signal-sequence carrying precursor proteins has been to shown to be significantly lower than for that of their mature, signal-sequence cleaved counterparts *(27,28)*. Thus, a translocation failure of mitochondrial and secretory proteins should generate conditions optimal for recognition and degradation by Ubr1p: availability of an uncleaved N-terminal sequence encoding an N-degron, accompanied by impaired or inhibited folding.

### Mis-translocated secretory and mitochondrial proteins are degraded by the N-end rule pathway

To test the above hypothesis, we examined ATP2Δ1,2,3, a translocation-defective mutant of Atp2p that is import-deficient and degraded in the cytosol *(29)*. ATP2Δ1,2,3 carries a Ubr1-QC-compatible P2 Val residue. Degradation was dependent on a combination of San1 and Ubr1 (**Figure 3A**). We substituted the P2 Val residue with a Ubr1-QC-*incompatible* Ser residue, and also mutated the P3 Leu to a Pro residue, producing ATP2Δ1,2,3-V2S,L3P. A P3 proline inhibits N-terminal acetylation, ensuring that any observed effect would not be due to blockage of the N-terminus *(21).* The degradation of ATP2Δ1,2,3-V2S,L3P was retarded when compared to that of ATP2Δ1,2,3 in Δ*san1* + UBR1 cells, while both forms of the substrate were stabilized to similar levels in Δ*san1*Δ*ubr1* cells, indicating that the decreased turnover rate of ATP2Δ1,2,3-V2S,L3P was the result of a reduction in Ubr1-mediated degradation (**Figure 3B**). Degradation of ATP2Δ1,2,3 was also dependent on Sse1, Ydj1, and Ssa1 and Ssa2 chaperones (**Figure S4**), hallmarks of UPS-dependent CytoQC and strong indicators of a misfolded substrate.

**Fig. 3.**
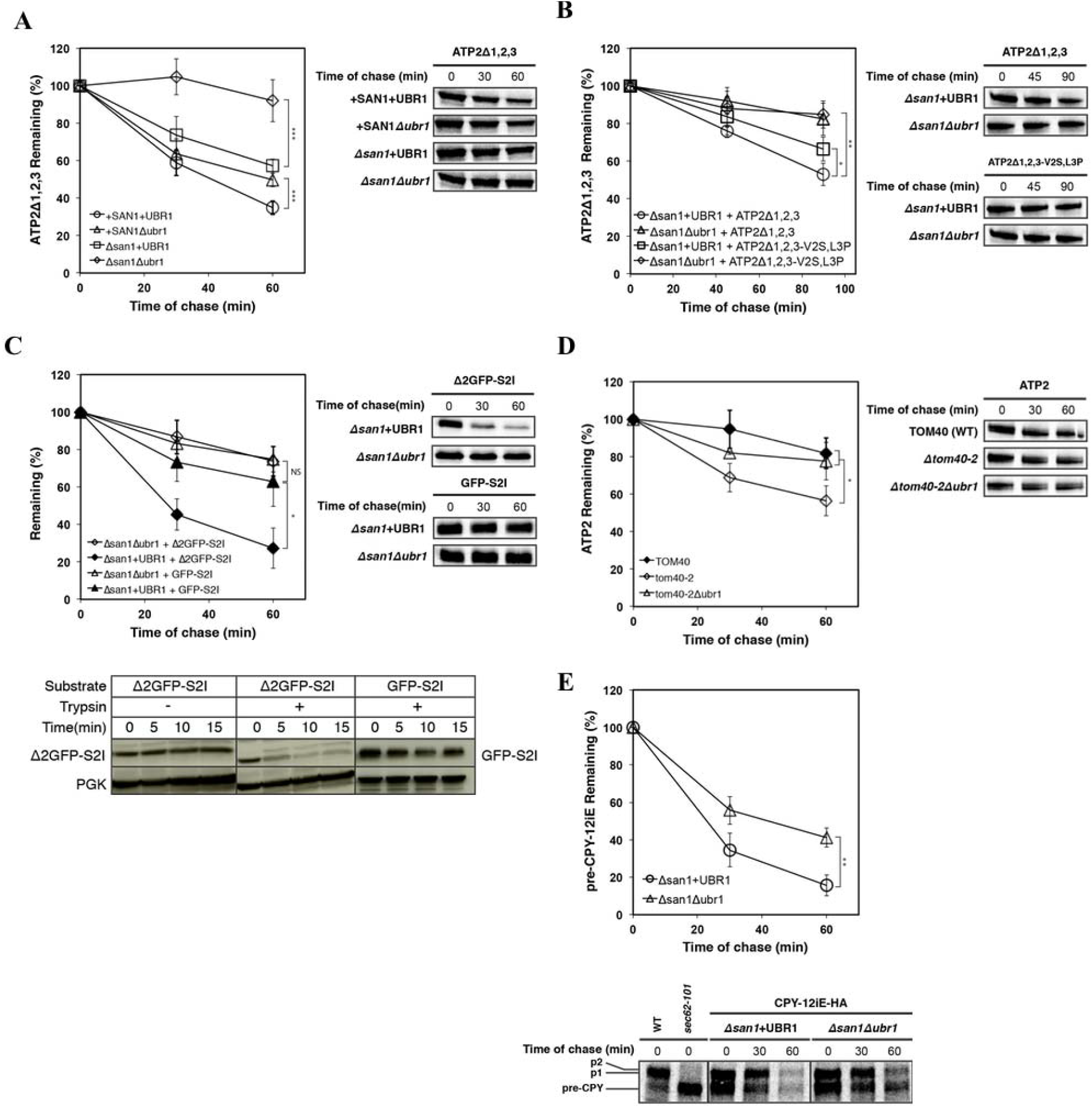
Mistranslocated forms of mitochondrial and secretory proteins are substrates of cytosolic quality control mediated by Ubr1 and the N-end rule pathway. (A) ATP2Δ1,2,3 turnover was determined by metabolic pulse-chase analysis in + SAN1 + UBR1 (wild-type), Δsan1 + UBR1, +SAN1Δubr1, and Δsan1Δubr1 strains (N = 5, biological replicates) (B) The turnover of ATP2Δ1,2,3 and ATP2Δ1,2,3-V2S,L3P was determined by metabolic pulse-chase analysis in Δsan1 + UBR1, and Δsan1Δubr1 strains (N = 4, biological replicates) (C) Turnover of GFP-S2I and Δ2GFP-S2I was compared by pulse-chase analysis in Δsan1Δubr and Δsan1 + UBR1 cells. Trypsin sensitivity assay (N = 1): postnuclear lysates were prepared from Δsan1Δubr cells and treated with 5.0μg/ml trypsin for the durations indicated. Protein was analyzed by immunoblotting with monoclonal anti-HA antibody. Endogenously expressed phosphoglycerate kinase (PGK) was assayed to serve as a folded protein control. (D) HA-tagged wild-type ATP2 protein expressed in TOM40 (wild-type), tom40-2, and tom40-2Δubr1 cells was analyzed by pulse-chase analysis. (E) Turnover of CPY-12iE-HA was analyzed by pulse-chase analysis in Δsan1 + UBR1 and Δsan1Δubr1 cells. (A-E) Metabolic pulse-chase: Cells were grown to log phase and shifted to 30°C for 30 minutes (experiment involving temperature sensitive *tom40-2* strains was performed at 37°C for all strains), followed by pulse-labeling with [35S]methionine/cysteine for 10 minutes and chased at the times indicated. Proteins were immunoprecipitated using anti-HA antibody and resolved by SDS- PAGE, then visualized by phosphoimager analysis. Samples from Δsan1Δubr1 cells expressing CPY-12iE-HA were exposed to a phosphor screen for an extended period (5 days vs 3 days for Δsan1 + UBR1 cells) for imaging and quantitative analysis due to a lower overall detection level of the expressed substrate compared to Δsan1 + UBR1 cells. Error bars, mean +/- SD of three independent experiments (N = 3, biological replicates) unless otherwise noted. Student’s t-test: *, p < 0.05; **, p < 0.01; ***, p < 0.005; Not Significant (NS), p > 0.5.

To assess the significance of a substrate’s misfoldedness to Ubr1-recognition of the substrate via a Ubr1-QC-compatible P2 residue, we examined wild-type GFP, which is expected to escape CytoQC-based degradation as it is a stable protein in yeast. The P2 Ser of GFP was substituted with Ile to generate GFP-S2I. GFP-S2I was significantly less susceptible to Ubr1 compared to Δ2GFP-S2I, which was degraded efficiently by Ubr1 (**Figure 3C**). Trypsin digestion confirmed the relative structural stability of GFP-S2I vs Δ2GFP-S2I (**Figure 3C**). These results indicate that degradation of a substrate via the N-end rule pathway is significantly enhanced when the substrate is also misfolded. This may be a result of the N-terminus of a folding-compromised substrate being aberrantly exposed, either spatially or temporally, making binding by Ubr1p binding more kinetically favorable. This is akin to the model of protein quality control proposed for misfolded proteins possessing Ac/N-end N-degrons *(1,17,30)*. Alternatively, it may be due to an increased availability of a misfolded substrate’s polyubiquitination-competent lysine residues, as one of the requirements of Ubr1-dependent degradation is access to a lysine in an unstructured region of the substrate *(12)*. Degradation of wild-type Atp2p in a temperature sensitive mitochondrial-import mutant, *tom40-2 (31)*, was also dependent on Ubr1 CytoQC (**Figure 3D**), suggesting that wild-type ATP2 is also misfolded when trapped in the cytosol. These results confirm that translocation-deficient Atp2p is an endogenous substrate of Ubr1-mediated CytoQC, and that its efficient degradation is dependent on its native Ubr1-QC-compatible P2 Val residue.

To determine if translocation-defective secretory pathway proteins are also subject to Ubr1 CytoQC, we generated an import-impaired mutant of vacuolar carboxypeptidase Y (CPY), CPY-12iE, in which a Glu residue is inserted at position 12 to disrupt the hydrophobic core of the native CPY signal sequence. Since any mature, vacuole-processed CPY-12iE would prevent accurate analysis of its pre-form levels, we utilized the fact that C-terminal processing of CPY-HA results in HA tag removal upon maturation (**Figure S5**). CPY encodes a Ubr1-QC-compatible P2 Lys. Degradation of pre-CPY-12iE significantly reduced by the absence of Ubr1p (**Figure 3E**, Δ*san1* + UBR1 vs Δ*san1*Δ*ubr1*), confirming that some subset of mis-translocated secretory proteins are also targets of Ubr1.

### P2 residues encode cellular location signals that mediate CytoQC of mistranslocated proteins

Studies have shown that mislocalized membrane proteins in the cytosol are targeted for degradation through exposed hydrophobic patches in both yeast and mammalian cells *(32–36)*. If mistranslocated proteins *without* Ubr1-QC-compatible P2 residues are primarily membrane proteins, the majority should be degradable through these pathways and rendered benign. To assess whether Ubr1 N-end rule based CytoQC, along with quality control pathways targeting hydrophobicity, could together provide an effective degradation system for clearing the cytosol of translocation-deficient, mis-localized proteins, we analyzed a set of 277 ORFs encoding signal sequence-containing proteins that was used in a previous study to determine the frequency of amino acids encoded at the P2 position in secretory proteins *(37)*. This set was filtered for duplicates and other inconsistencies, resulting in 273 proteins which were then categorized as soluble or membrane proteins based on existing literature. If existing literature supporting soluble or membranous topology for a particular ORF did not exist, data from SignalP, Kyte-Doolitle hydrophilicity profiling, and Phobius, a transmembrane domain prediction tool, were together utilized for the determination of topology (refer to Materials and Methods). Strikingly, 87.5% (49/56) of ER proteins encoded with Ubr1-QC-*incompatible* residues are membrane proteins, strongly differing from the full set of signal-sequence bearing proteins, for which only 64.1% (173/270) are membrane proteins (p < 0.001, *x*^2^ = 11.7) (**Figure 4A; Tables S12-S16**). An even larger difference was observed when compared to the relative frequency of membrane proteins in the set of Ubr1-QC-compatible P2-residue-encoded proteins (57.9% (124/214); p < 5 × 10^−5^, *x*^2^ = 16.8). These results indicate that the vast majority of ER proteins which are not compatible with Ubr1 are also, at a much higher relative frequency, membrane proteins, which is expected to facilitate their degradation through pathways that target exposed hydrophobicity, such as San1-mediated CytoQC.

**Fig. 4.**
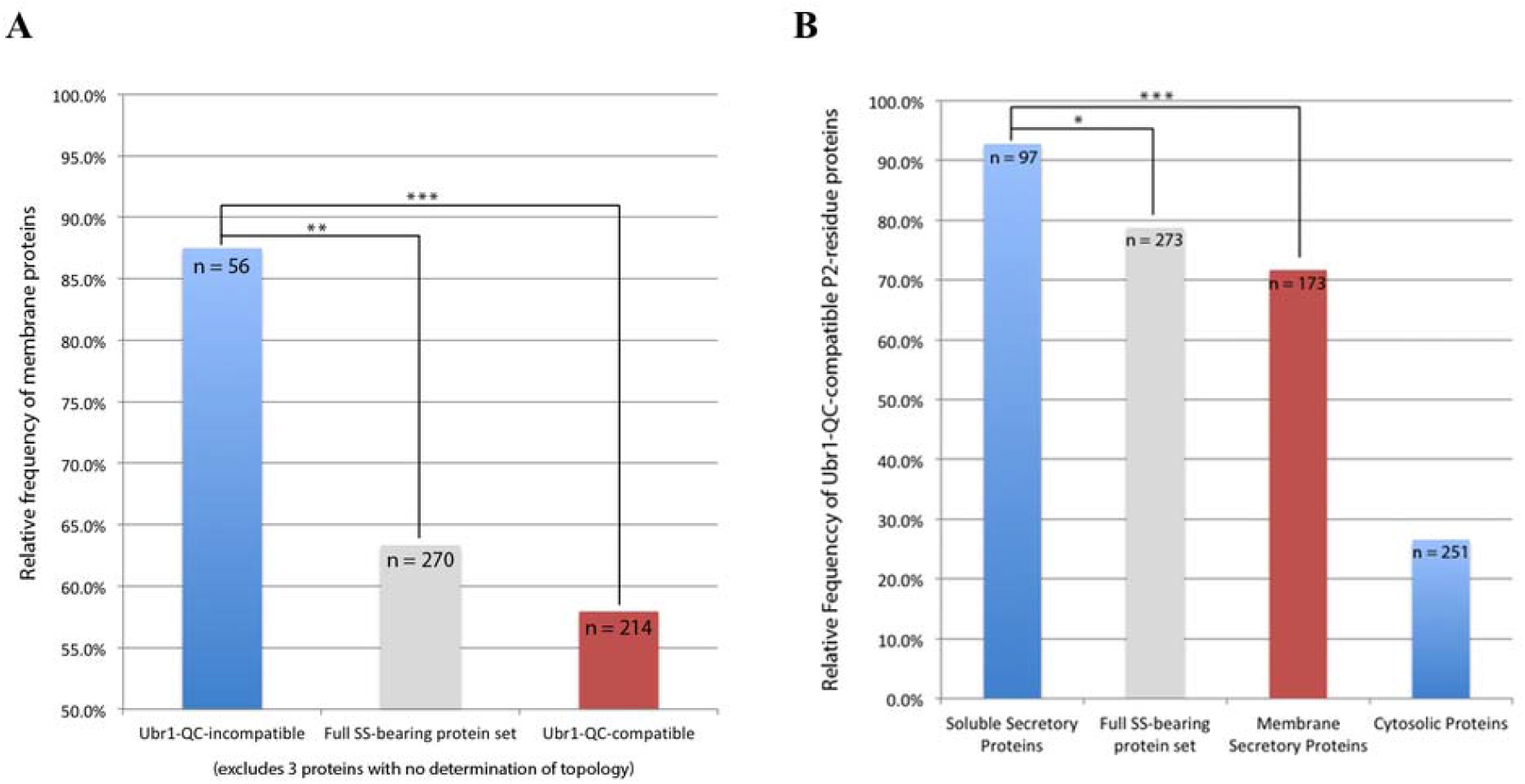
The majority of secretory proteins encoding Ubr1-QC-*incompatible* P2-residues are membrane proteins, while the majority of secretory proteins encode Ubr1-QC-compatible P2 residues. (A) The percentages of proteins that are membrane-bound was calculated for three categories derived from the full set of 270 signal-sequence bearing proteins for which topology was determined: those encoded with Ubr1-QC-compatible P2 residues, those encoded with Ubr1-QC-*incompatible* P2 residues, and the full set (see Materials and Methods; Tables S12, S14A, S15A, S16). Chi-square test: Ubr1-incompatible (N = 56, proteins) vs full set (N = 270, proteins): **, p < 0.001, *x*^2^ = 11.7, *df* = 1; Ubr1-incompatible (N = 56, proteins) vs Ubr1-compatible (N = 214, proteins): ***, p < 5 × 10^−5^, *x*^2^ = 16.8, *df* = 1. (B) The percentages of proteins encoded with Ubr1-QC-compatible P2 residues was calculated for three categories of proteins derived from the full set of 273 signal-sequence bearing proteins: those determined to be soluble, membranous, and the full set (see Materials and Methods; Tables S12, S14A, S15A, S16). Chi-square test: Soluble (N = 97, proteins) vs full set(N = 273, proteins): *, p < 0.005, *x*^2^ = 9.7, *df* = 1. Soluble(N = 97, proteins) vs membrane set(N = 173, proteins): ***, p < 5 × 10^−5^, *x*^2^ = 16.8, *df* = 1. Cytosolic protein set (N = 251, proteins) from Forte et al., 2011 *(39)*.

Interestingly, statistically significant differences were seen in the relative frequency of proteins encoding Ubr1-QC-compatible P2 residues when comparing the set of soluble proteins with membrane proteins (Chi-square test: p < 5 × 10^−5^, *x*^2^ = 16.8), as well as with the full set of signal-sequence bearing proteins (p < 0.005, *x*^2^ = 9.7): 92.8% (90/97) of the soluble protein set possess Ubr1-QC-compatible P2 residues, versus 78.8% (215/273) of the full set of signal-sequence bearing proteins, and only 71.7% (124/173) of the membrane proteins. (**Figure 4B; Table S12-S16**). Thus, while the majority of signal-sequence bearing proteins are susceptible to Ubr1, soluble secretory proteins in particular are primed for degradation through this pathway. In contrast, only ∼26.6% of cytosolic proteins possess Ubr1-QC-compatible P2 residues based on an analysis of a representative set of 251 randomly selected cytosolic proteins *(37)* (**Figure 4B**; **Table S11)**. This minority of cytosolic proteins with Ubr1-QC-compatible P2 residues should fold efficiently in their native folding environment, thereby escaping targeting by Ubr1 CytoQC, as demonstrated in stability experiments involving folded vs misfolded GFP (**Figure 3C**). Why these cytosolic proteins have also evolved to encode N-degrons would be an interesting avenue of study.

## Discussion

The findings presented here broaden our understanding of how eukaryotic cells ensure the cytosol is kept in a healthy state mostly clear of foreign actors that could disrupt normal cellular processes such as signaling, protein synthesis, and trafficking. Failures in SRP-mediated ER targeting causes the mistargeting of secretory proteins to the mitochondria, resulting in mitochondrial dysfunction *(38)*, while the depletion of nascent polypeptide-associated complex (NAC) results in the incorrect import of mitochondrial proteins into the ER lumen*(39)*. Such aberrant cross-organelle mis-targeting would also be mitigated by the Ubr1 P2-dependent CytoQC pathway, capturing such proteins in the cytosol before mis-targeting is able to occur. We propose that there is an intrinsic pressure for secretory and mitochondrial proteins to be encoded with Ubr1-QC-compatible P2 residues so they can be efficiently degraded in the event of mislocalization as a result of translocation failure. Soluble ER proteins are particularly targetable by Ubr1 since the vast majority possess Ubr1-QC-compatible P2 residues. That they would not be recognized by hydrophobicity-based degradation pathways due to their lack of transmembrane domains, and possession of only moderately hydrophobic signal sequences, necessitates Ubr1-QC-compatibility to be efficiently cleared from the cytosol. Together, our findings also suggest that P2 residues are utilized as *de facto* cellular location signals which help to ensure the fidelity of protein localization. This system of P2-encoded location signaling facilitates N-end rule pathway-mediated degradation of a majority of mis-translocated secretory and mitochondrial proteins in the cytosol.

The discovery of Met-Φ Arg/N-end degrons by the Varshavsky and Hwang labs *(15)* was surprising for several reasons. Firstly, N-terminal Met was originally classified as a stabilizing residue. Secondly, the existence of Met-Φ sequences as N-degrons meant that a large number of proteins translated in the cytosol may be susceptible to recognition by the Arg/N-end rule pathway. Our characterization of Met-Φ N-degrons as part of an expanded set of Arg/N-end rule N-degrons that are recognized when possessed by folding-impaired, mistranslocated proteins, helps to identify a specific common physiological occurence under which these N-degrons take effect: translocation failure. Our findings thus bring to light large swaths of secretory and mitochondrial proteins that are degradable via the Arg/N-end rule pathway. Importantly, they do not require an endoproteolytic cleavage or N-terminal modification to expose destabilizing residues, as the case with the majority of previously characterized substrates in Arg/N-end rule pathway-dependent mechanisms *(12)*. Instead, they rely on the qualities of misfoldedness and mislocalization to enable the utilization of their natural N-terminal N-degrons as effective degradation signals.

Confoundingly, the localization of Ubr1 is predominantly in the nucleus, where some of its misfolded substrates are imported and ubiquitinated by it *(40)*. A smaller fraction of Ubr1 also operates on substrates in the cytosol itself *(8,16,41)*. Whether the majority of mislocalized P2-residue-dependent Ubr1 substrates are engaged by the cytosolic form of Ubr1, or are first trafficked to the nucleus and subsequently recognized by the nuclear form of Ubr1, remains to be determined. However, since we observed an even lower usage frequency of Ubr1-QC-compatible P2 residues in the case of nuclear-localized proteins as compared to cytosolically-localized proteins, Ubr1 presence and degradation activity in both nuclear and cytosolic compartments is still congruent with our model of P2-based location signaling and degradation (**Figure 2F**).

Herpes simplex virus 1-encoded microRNA has been shown promote the accumulation of β-amyloid through the inhibition of Ubr1 activity, while aggregation-prone fragments of neurodegeneration-associated TDP43, Tau, and α-synuclein, are substrates of the N-end rule pathway *(42,43)*. An intriguing avenue of research would be to investigate if the P2-residue dependent degradation of translocation-deficient proteins by Ubr1 described here is mirrored in mammalian cells, and if so, whether disruption of the pathway is involved in the accumulation of such detrimental disease factors.

An additional surprising corollary to our discovery is that the N-end rule pathway, through the recognition of P2-encoded cellular location signals, plays a primary role in enforcing the compartmentalization of secretory and mitochondrial proteins that would otherwise adversely affect critical cellular activities in the cytosol. It is widely accepted that mitochondria and plastids in eukaryotes evolved from endosymbiotic partnerships, whereby host prokaryotic cells engulfed ancient aerobic bacteria and gained the benefit of increased energy production, while the new cellular tenants were consequently shielded from the environment *(44)*. Over time, gene transfer from endosymbionts to the host cell genome occurred that conferred physiological fitness benefits, and evolved further into complex systems that utilized transit peptides and specialized import machinery to direct host-encoded proteins in the cytosol back to the endosymbiont *(45)*. However, how did early eukaryotes deal with endosymbiont-derived proteins that could not make it to their intended destination? Based on the results presented here, we propose that the N-end rule pathway played a critical role in the evolution of advanced eukaryotic cells by providing a mechanism to destroy proteins intended for endosymbiotic organelles that failed to successfully transit. It is thus no surprise that the N-terminal signal sequences of ER and mitochondrial proteins largely coincide with the set of P2 residues that are Ubr1-compatible, with soluble translocated proteins in particular exhibiting nearly total concordance based on our analyses of unbiased datasets. In this way, the N-end rule pathway may have been the key enabler for eukaryotic cells to fully exploit the benefits of harboring endosymbiotic organelles by compensating for the negative physiological effects associated with mistargeted proteins.

## Acknowledgments

I thank my ex-graduate advisor Davis Ng for his expert guidance and exchange of ideas throughout this work. I am grateful to Nassira Bedford for assistance with several unpublished experimental studies, and Chengchao Xu and Eric Fredrickson for their critical review of the manuscript and invaluable feedback and advice.

## Author contributions

AT designed the studies, performed the experiments, and wrote the manuscript.

## Data and materials availability

All data is available in the main text or the supplementary materials. Other raw data, code, gel scans, etc., are available upon request from the author.

## Supplementary Information

### Materials and Methods

Note: All non-bioinformatic biological experiments reported here were part of thesis work performed by Anthony Tran at National University of Singapore for his doctoral dissertation in the lab of Associate Professor Davis Ng at the National University of Singapore [46].

### Study Design

#### Research objectives and units of investigation

The over-arching goal of the project was to systematically determine the effects that altering N-terminal sequences had on the degradation of misfolded proteins by Ubr1 and the N-end rule pathway. Specifically, this was done by selecting a known cytosolic quality control substrate, Ste6*C, creating a full set of P2-residue variants derived from it, and assessing degradation rates of the substrates in yeast cell lines that differred in the presence or absence of Ubr1 and San1, the E3 ligases of interest. The P2 position was selected for analysis as it is the primary determinant in the N-terminal processing of proteins and the resultant N-terminal sequence of a protein prior to downstream processing of signal sequences [18-19]. Based on these initial degradation assays, global bioinformatic analysis was used to determine the cellular compartments that had the highest frequency of proteins with Ubr1-QC compatible P2 residues encoded. The finding that mitochondrial and secretory proteins encode Ubr1-QC-compatible P2 residues more frequently led us to formulate the follow-up hypothesis that they are the primary targets of Ubr1 quality control in the cytosol in the event of translocation failure. This hypothesis then drove us to perform the targeted in-depth bioinformatic and biochemical analysis of representative secretory and mitochondrial protein sets and substrates in relevant cell-lines and translocation conditions to prove the validity of the hypothesis, finally leading to formulation of the new model for Ubr1-mediated CytoQC.

#### Replicate and selection of end-points

Pulse-chase experiments, used to determine the rates of degradation of endogenous and engineered substrates in various cell backgrounds, were performed in sets of at least 3 biological replicates unless otherwise noted. Three replicates is an accepted standard minimum for many biological experiments, and is also widely utilized in previously published reports which employed the same protocol. Furthermore, it allowed us to determine means, standard deviations, and verify statistically significant differences (or lack of differences) between strains and/or substrates. Intermediate end-points for pulse-chase assays were selected either based on those used in previous studies analyzing the same substrate type, or through pilot experiments to determine durations that would allow discernable, statisically significant deviations to surface in relevant strains. Replicates were generally performed on separate days and times to account for possible fluctuations in laboratory environment and equipment performance. Strains expressing analyte substrates were freshly inoculated for each replicate experiment to account for normal or unexpected variations in growth environment, growth medium, and biological activity.

#### Experimental Design

Biological experiments were conducted in controlled laboratory environments with specific incubation temperatures, times, treatment quantities, durations, and measurement methods, as described below in the sections applicable for each experiment.

#### Sample sizes

For bioinformatic analyses assessing the relative frequencies of P2 residue amino acid usage in the yeast proteome based on cellular localization and ORF sequences, a widely-cited GFP-fusion localization study [22] was utilized in order to determine and compare P2 amino acid residue encoding frequency for proteins in different cellular compartments. For the best assessments of this parameter, the largest sample size available from the study for each cellular compartment analyzed was utilized, and is described in more detail in the *Bioinformatic Analysis* section.

For targeted bioinformatic analyses of P2 amino acid usage and topological determination of secretory pathway proteins based on a combination of literature review and Kyte Doolittle/Phobius methods, we utilized a published set of 277 signal-sequence bearing proteins/ORFS that was previously used for determining P2 residue frequency for proteins in the secretory pathway [39]. Using a previously vetted and published secretory protein set eliminated chance of bias for the purpose of our analysis of P2 Ubr1-QC-compatibility. Based on the same logic, the set of 251 randomly selected cytosolic proteins from the same study [39] was used for our analysis of P2 amino acid residue frequencies in the cytosol to prevent potential bias in selecting cytosolic proteins for our analysis.

### Strain and Antibodies

Yeast strains used in this study are listed in Table S28. Anti-HA monoclonal antibody (HA.11) was sourced from Covance (Princeton, New Jersey). Monoclonal anti-3-phosphoglycerate kinase (PGK) was sourced from Invitrogen (Carslbad, California). Anti-CPY antibody was a gift from Reid Gilmore (University of Massachusetts, Worcester, MA).

### Plasmids and Primers

Standard cloning procedures were utilized for the construction of plasmids [47]. Unless otherwise stated, exogenously expressed substrates possess an engineered single hemagglutinin (HA) epitope tag attached to the C-terminus. Ste6*C, Δ2GFP, ATP2Δ1,2,3, and their derivatives, were expressed under control of a high expression, constitutive TDH3 (glyceraldehyde-3-phosphate dehydrogenase) promoter in yeast centromeric plasmids. HA-tagged Prc1p (CPY/carboxypeptidase Y) and its derivatives were placed under control of its native constitutive endogenous promoter in yeast centromeric plasmids. Site-directed mutagenesis of the original constructs expressing Ste6*C, Δ2GFP, and ATP2Δ1,2,3 was performed to generate mutant substrates.

pAT1: A fragment carrying the PRC1 promoter was PCR amplified from pSW119 with BamHI and NotI restriction ends. The amplified fragment and pSW119 were digested with NotI and BamHI and ligated to generate pAT1.

pAT32: A fragment encoding the TDH3 promoter, followed by Ste6*C-HA, followed by the ACT1 terminator sequence, was PCR amplified from pRP22 with primers AT270 and AT273 and digested with NotI and XhoI. The fragment was then ligated into an empty pRS316 vector to generate pAT32.

pAT33-pAT51: pAT33 through pAT51 (expressing Ste6*C-I2K, Ste6*C-I2Y, Ste6*C-I2F, Ste6*C-I2A, Ste6*C-I2L, Ste6*C-I2E, Ste6*C-I2V, Ste6*C-I2G, Ste6*C-I2R, Ste6*C-I2M, Ste6*C-I2P, Ste6*C-I2W, Ste6*C-I2N, Ste6*C-I2D, Ste6*C-I2H, Ste6*C-I2Q, Ste6*C-I2C, Ste6*C-I2T, Ste6*C-I2S) were constructed by mutation of the base-pairs encoding the 2nd residue of Ste6*C through site-directed mutagenesis using primers AT21-AT39 and pRP22 as a template.

pAT52: A fragment encoding residues 1201-1290 of the STE6 ORF followed by a hemagglutinin epitope (HA-tag) sequence was PCR amplified from yeast genomic DNA using primers AT40 and AT41. The fragment was digested with BamHI and XbaI and ligated to pRP22 digested with BamHI and XbaI generating pAT52.

pAT55-pAT57: pAT55 through pAT57 (expressing Δ2GFP-S2I, Δ2GFP-S2F, Δ2GFP-S2K) were constructed by mutation of the base-pairs encoding the 2nd residue of Δ2GFP through site-directed mutagenesis using primers AT18-AT20 and pRP44 as a template.

pAT61: A 741bp fragment of the GFP ORF followed by the hemagglutinin (HA) tag sequence was PCR amplified using mutational primers AT189 and AT190 and pAT7 as a template. The 20 resultant PCR product carrying the GFP ORF with the 2nd residue mutated to an isoleucine was digested with BamHI and XbaI and ligated to pAT7 digested with BamHI and XbaI generating pAT61.

pAT64: A 1563-bp fragment carrying the ATP2 ORF followed by the hemagglutinin epitope (HA-tag) sequence was PCR amplified from yeast genomic DNA using primers AT244 and AT226. The fragment was digested with BglII and XbaI and ligated to pAT7 digested with BamHI and XbaI generating pAT64.

pAT65: pAT66 was digested with ClaI and XhoI to release a 2671-bp fragment encoding the TDH3 promoter, ATP2Δ1,2,3-HA, and ACT1 terminator sequences. This fragment was ligated into an empty pRS316 vector digested with ClaI and XhoI to generate pAT65.

pAT66: A 1506-bp fragment carrying the ATP2 ORF with deletions of residues 5-12, 16-19, and 28-34, followed by the hemagglutinin epitope (HA-tag) sequence was PCR amplified from yeast genomic DNA using primers AT246 and AT226. The fragment was digested with BglII and XbaI and ligated to pAT7 digested with BamHI and XbaI generating pAT66.

pAT68: pAT68 (expressing ATP2Δ1,2,3-V2S,L3P-HA) was constructed by mutation of the sequences encoding the 2nd and 3rd residues of ATP2Δ1,2,3-HA, from valine to serine at the P2 position, and leucine to proline at the 3rd position, through site-directed mutagenesis using primer AT275 and pAT66 as a template.

pAT72: pAT72 (expressing PRC1-12iE) was constructed by the insertion of a three base-pair sequence encoding a glutamic acid residue at the 12th codon position of the PRC1 ORF through site-directed mutagenesis using primer AT280 and pXW92 as a template.

### Metabolic Pulse-Chase Assay

Yeast cells were grown to log phase at 30°C (25°C for temperature sensitive strains). 3 OD600 units of cells were resuspended in 0.9 ml of SC or SC selective media and incubated at 30°C (37°C for temperature sensitive strains) for 30 minutes. Pulse labeling was then initiated with the addition of 82.5 μCi of [35S]Met/Cys (EasyTagTM EXPRESS 35S, PerkinElmer) for 5 or 10 minutes depending on the labeling efficiency of the substrate of interest. Label was chased with the addition of excess cold methionine and cysteine to a final concentration of 2 mM. At the appropriate timepoints, pulse labeling/chase was terminated by the addition of 100% TCA to a final concentration of 10%. Immunoprecipitation of samples and resolution by SDS-PAGE were carried out as described [48]. Phosphor screens exposed to gels (1 to 5 day exposure, depending on substrate expression level) were scanned with a Typhoon^TM^ phosphoimager and the visualized bands of interest quantified using ImageQuant TL software (GE Healthcare Life Sciences, Uppsala, Sweden) or ImageJ. Background signal from screen exposure was subtracted. All results presented are the mean ± SD of three independent experiments (N = 3, biological replicates) unless otherwise specified.

### Trypsin Sensitivity Assay

Yeast cells expressing the substrate of interest were grown to log phase (0.4-0.6 OD) and resuspended in cytosol buffer (20 mM HEPES-KOH, pH 7.4, 14% glycerol, 100 mM KOAc, and 2 mM MgOAc) at a concentration of 20 OD/mL. 1 mL of this resuspension was transferred to a 2 ml screw-cap tube and homogenized by vortexing for 30 seconds in the presence of 1 ml of 0.5 mm diameter zirconium beads followed by a 1 minute incubation at 4°C. This was performed for 5 cycles. The homogenate was transferred to 1.5 ml eppendorf tubes. 0.6 ml of fresh cytosol buffer was used to wash the beads and pooled with the original homogenate. Post-nuclear lysate was isolated by pelleting at 500xg for 5 minutes and transferring the supernatant to a fresh tube. The post-nuclear lysate was incubated at 30°C for 5 minutes, followed by the addition of trypsin to a concentration of 5μg/ml. Samples were vortexed and incubated for 30°C, with 100ul aliquots taken at the indicated timepoints and mixed with 11.1ul of 100% TCA in fresh 1.5 ml eppendorf tubes. Aliquots were kept on ice for 5 minutes and pelleted at 14000 rpm for 20 minutes at 4°C. Supernatant was discarded, sample pelleted again briefly, and supernatant again discarded. The pellet was resuspended in 10ul of TCA resuspension buffer (100 mM Tris-HCl pH 11.0, 3% SDS, 1 mM PMSF) by cycles of boiling at 100°C and vortexing. Samples were pelleted at 4°C to remove SDS and other insoluble particles, and the resultant supernatant transferred to a fresh tube. Analysis by SDS-PAGE/Western Blotting was performed using the appropriate antibodies.

### Edman Degradation N-terminal Seqeuncing

Yeast cells expressing the protein of interest were grown to 1 OD/mL in selective media. 800OD of yeast cells were harvested at 3000xg for 15 minutes, washed once with 1x PBS, and pelleted again at 3000xg. Cells were washed in IP/NP40/PIC/DTT (50 mM Tris-HCl, pH8 150 mM NaCl, 0.1% NP-40, 0.1 mM DTT), pelleted at 3000xg, and resuspended in IP/NP40/DTT containing protease inhibitors (complete, mini, EDTA-free protease inhibitor cocktail tablet, Roche) at a concentration of 50 OD/ml. 1 mL aliquots of the resuspension were transferred to a 2 ml screw-cap tubes and homogenized by beadbeating for 30 seconds using a Mini-BeadBeater cell disrupter (Biospec Products) followed by a 5 minute incubation on ice; beadbeating and incubation on ice was repeated for 6 cycles in the presence of 1 ml of 0.5 mm diameter zirconium beads. The homogenate was transferred to 1.5 ml eppendorf tube. 0.6 ml of fresh cytosol buffer was used to wash the beads and pooled with the original homogenate. Lysate was cleared by centrifugation at 14000 rpm and transferred to a fresh tube. Lysate was incubated with 65uL of Roche Anti-HA affinity matrix per 4 mL of lysate for 2 hours or overnight at 4°C. Affinity matrix was spun down at 2700 rpm for 1 minute and washed with ice cold IP/NP40/PIC/DTT three times, followed by one wash with cold IP buffer to remove residual NP40. Bound proteins were eluted from matrix through the addition of protein loading buffer (PLB) and subsequent boiling at 100°C for 10 minutes. Immunoprecipitated proteins were resolved on 12% SDS-PAGE gels and transferred to a PVDF membrane. Membrane was washed with dIH2O for 1 minute (3 × 1 minute, shaking at 70 rpm) to eliminate traces of SDS, Tris, glycine, and other reagents that have the potential to interfere with Edman chemistry. The membrane was then stained with Coomassie Brilliant Blue (0.1% CBB, 5% acetic acid, 50% methanol) for 5 minutes by shaking at 70 rpm. Membrane was quickly destained with 50% methanol (3 × 1 minute, shaking at 70 rpm). Band containing the protein of interest was excised from the membrane and cut into smaller pieces to facilitate sample analysis. Membrane fragments were loaded into an ABI Procise 494 Sequencer for sequencing using standard manufacturer recommended protocols.

### Bioinformatic Analysis

The raw data file containing protein translations for systematically-named ORFs was obtained from Saccharomyces Genome Databse (SGD). To facilitate parsing with PHP, quotation marks in protein descriptions were removed, and a termination character (“@”) was added to the end of the file. (http://downloads.yeastgenome.org/sequence/S288C_reference/orf_protein/; orf_trans.fasta.gz) A PHP script was written to extract from this data file the systematic name of each ORF and the first 2 residues in its respective protein sequence, and subsequently output into an SQL database; a list of 5887 ORFS and their N-terminal sequence were produced (Table S1). The list of proteins exclusively localized to each of the main protein localization categories (nuclear, mitochondrial, cytoplasmic, or secretory) was determined based on complete or partial localization to that category, as determined by the localization terms assigned to each protein through a GFP-fusion localization method [22]. Nuclear proteins included those assigned the following localization terms: *nucleus, nucleolous*, and *nuclear periphery*. Secretory proteins included those assigned the following terms: *ER, Golgi, vacuole, endosome*, and *peroxisome*. Proteins assigned the term *mitochondrion* were categorized as mitochondrial. Proteins assigned with the term *cytoplasm* were categorized as cytosolic. To determine the relative total abundance of proteins with nuclear, cytosolic, secretory, or mitochondrial localization that encode Ubr1-QC-compatible or Ubr1-QC-incompatible P2 residues, we summed the abundance levels of proteins in each subset as quantified in the GFP-localization study, and divided it by the aggregate abundance of all proteins within the full localization category (Table S17-S27).

#### Categorization of signal-sequence bearing proteins

A published set of 277 signal-sequencing bearing proteins was analyzed [39] (Table S12). Two of this original set were duplicate entries (AIM6/YDL237W duplicate of LRC1, FLO11/YIR019C duplicate of MUC1) and were not included in the analysis. Two were not present in the YeastSGD database and were also excluded (YCR012C, and YJL052C) (Table S13). After these exclusions, a manual review was performed on each protein to identify literature that supported soluble or membranous topology. 211 of the proteins were successfully categorized through the review of literature (Tables S14-S16, S14A, S15A). For 59 proteins for which supporting literature could not be found, protein sequences of each were analyzed with SignalP 4.1 using the default optimized parameters. Proteins that were not predicted to possess a signal sequence cleavage site were categorized as membrane proteins. To assess the topology of proteins that were predicted to possess a signal sequence cleavage site, Kyte-Doolittle hydropathy profiles were obtained, and membrane prediction with Phobius performed. Kyte-Doolittle analysis was performed with a window size of 19. Proteins with a hydrophobicity score of greater than 1.8 after the predicted cleavage site were categorized as membrane proteins. Proteins that did not have a hydrophobicity score of greater than 1.8 after the predicted cleavage site were categorized as soluble proteins. 3 proteins which did not have consensus between Kyte-Doolittle and Phobius analysis were excluded from relative frequency calculations involving topology.

### Statistical Analysis

A standard of 3 replicates for biological experiments was performed in the determination of means, standard deviations, and statistical significance, unless otherwise noted. Unpaired, two-tailed Student’s t-test was performed to assess statistical significances in differences in degradation rates based on pulse-chase experiments.

Chi-square analysis was performed to assess statistically signicant differences in P2-residue amino acid frequency distributions between protein sets across all 20 possible residues using 19 degrees of freedom. Pair-wise chi-square analysis was performed when comparing the relative frequency of a single residue between two protein sets using 1 degree of freedom. Pair-wise chi-square analysis was performed when comparing the relative frequency of Ubr1-QC-compatible proteins between two protein sets, and when comparing the relative frequency of membrane proteins between two protein sets, using 1 degree of freedom.

Detailed N values (either biological experimental replicates, or samples) are included in the respective Fig legends.

P-value thresholds were set to 0.05 or less, and are indicated in the respective Fig legends as asterisks between the relevant sample data, or multiple asterisks in the case of lower p-value thresholds.

**Fig. S1.**
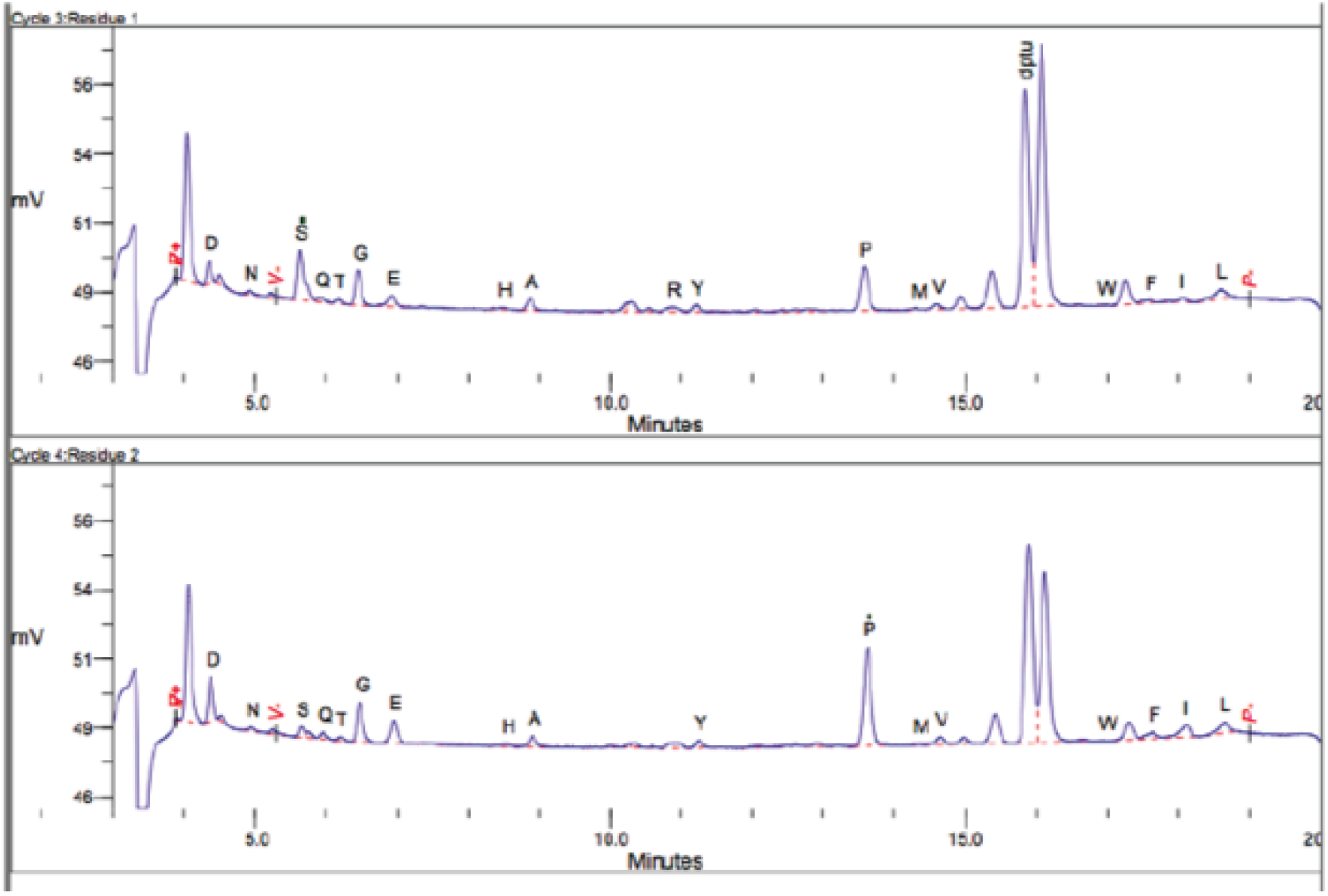
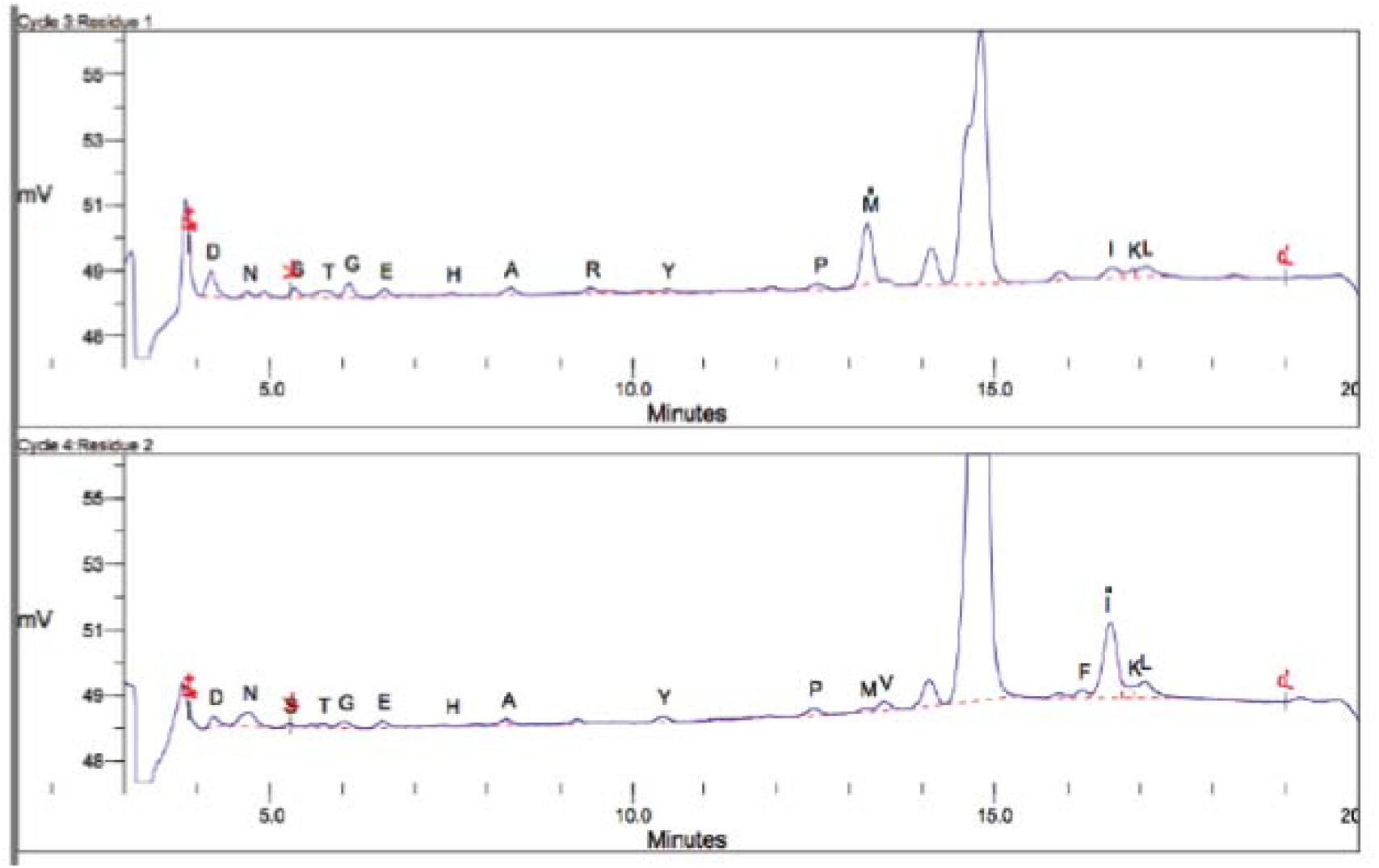
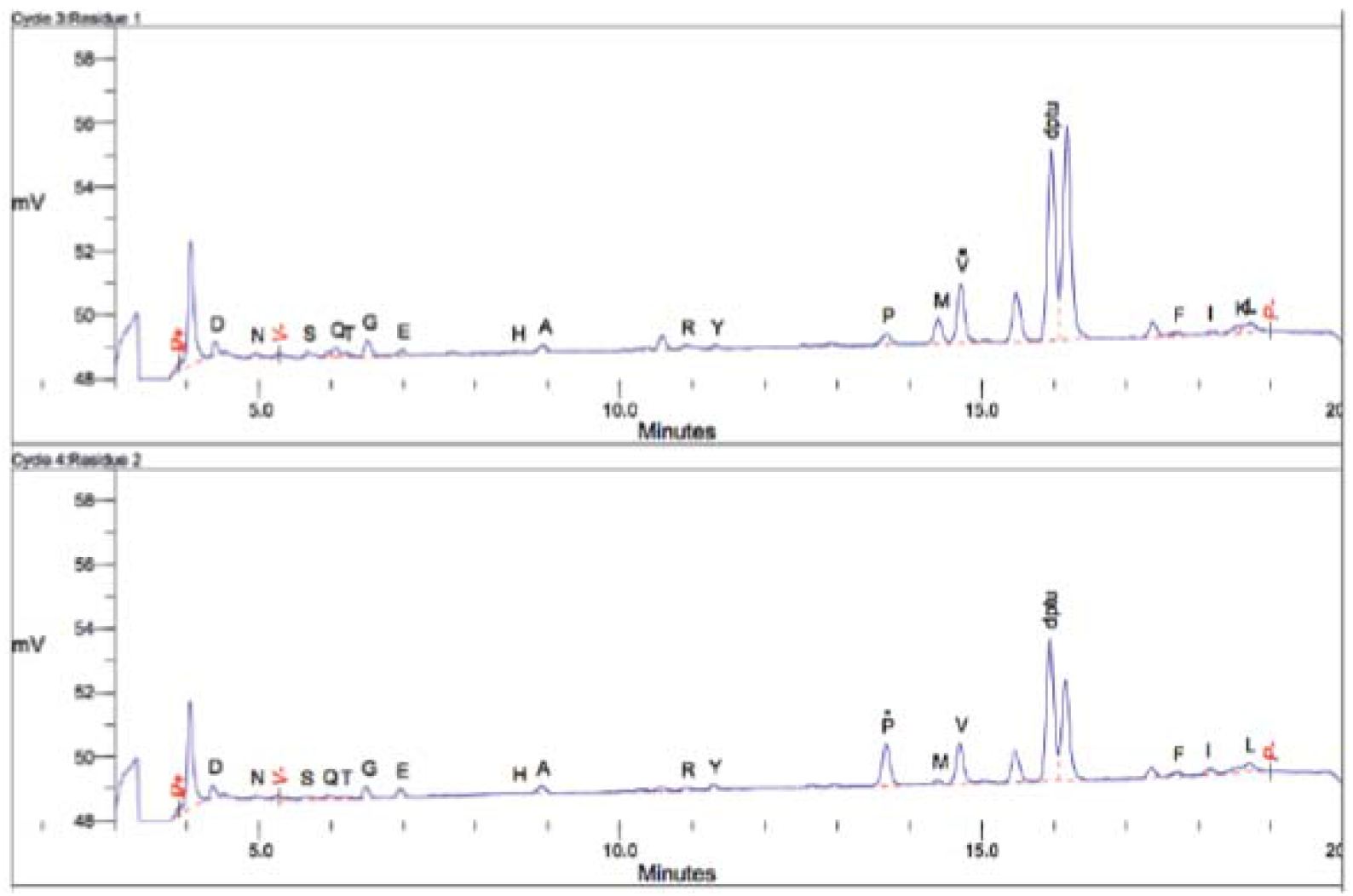
N-terminal sequencing of Ste6*C constructs. (A) Lysate was prepared from m Δsan1Δubr1 cells expressing Ste6*C-HA. Protein was immunoprecipitated from lysate with anti--HA affinity matrix (Roche), resolved by SDS-PAGE, and transferred to PVDF membrane.e. Protein band carrying Ste6*C-HA was excised from the membrane and sequenced via Edman an degradation. (B) Lysate was prepared from Δsan1Δubr1 cells expressing Ste6*C-HA-I2S.S. Protein was immunoprecipitated from lysate and sequenced as described in Fig S1A. (C) Lysate te was prepared from Δsan1Δubr1 cells expressing Ste6*C-HA-I2 V. Protein was as immunoprecipitated from lysate and sequenced as described in Fig S1A.

**Fig. S2.**
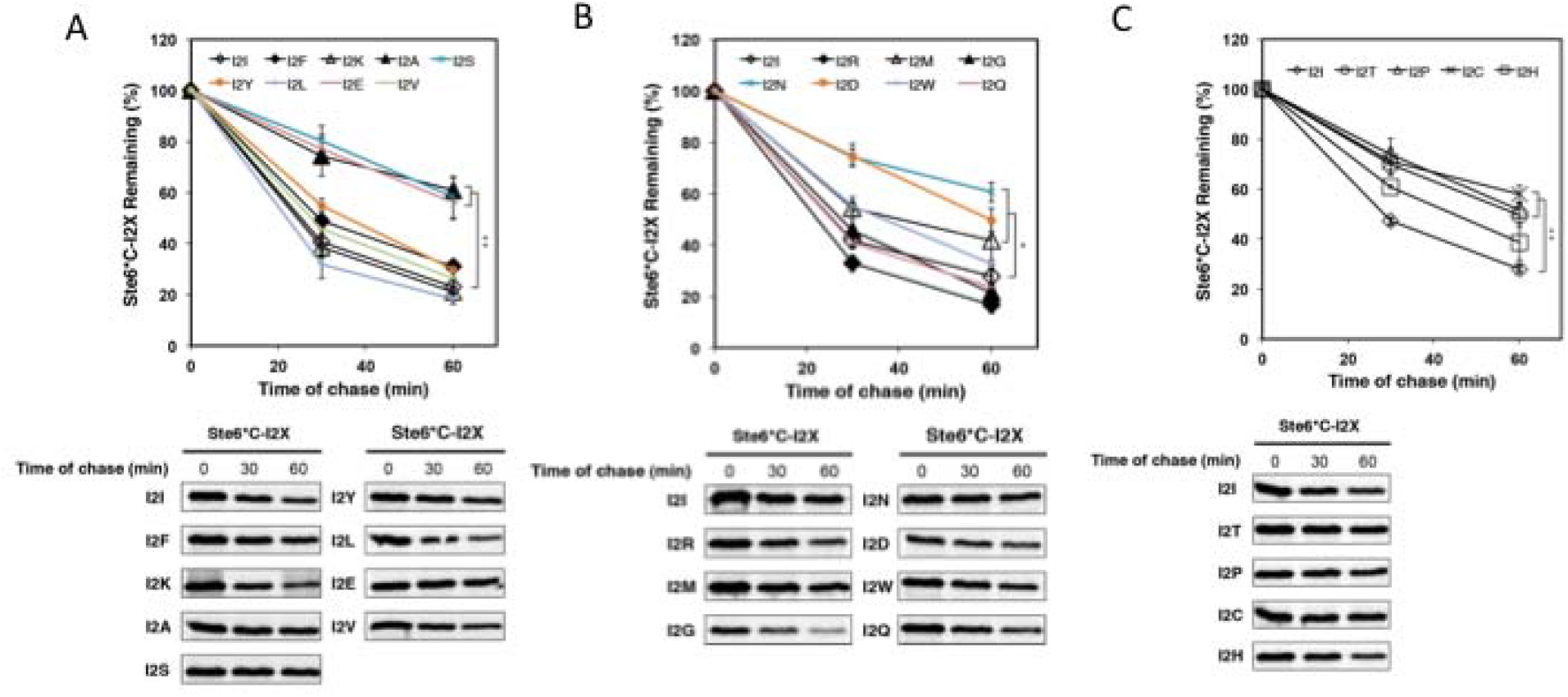
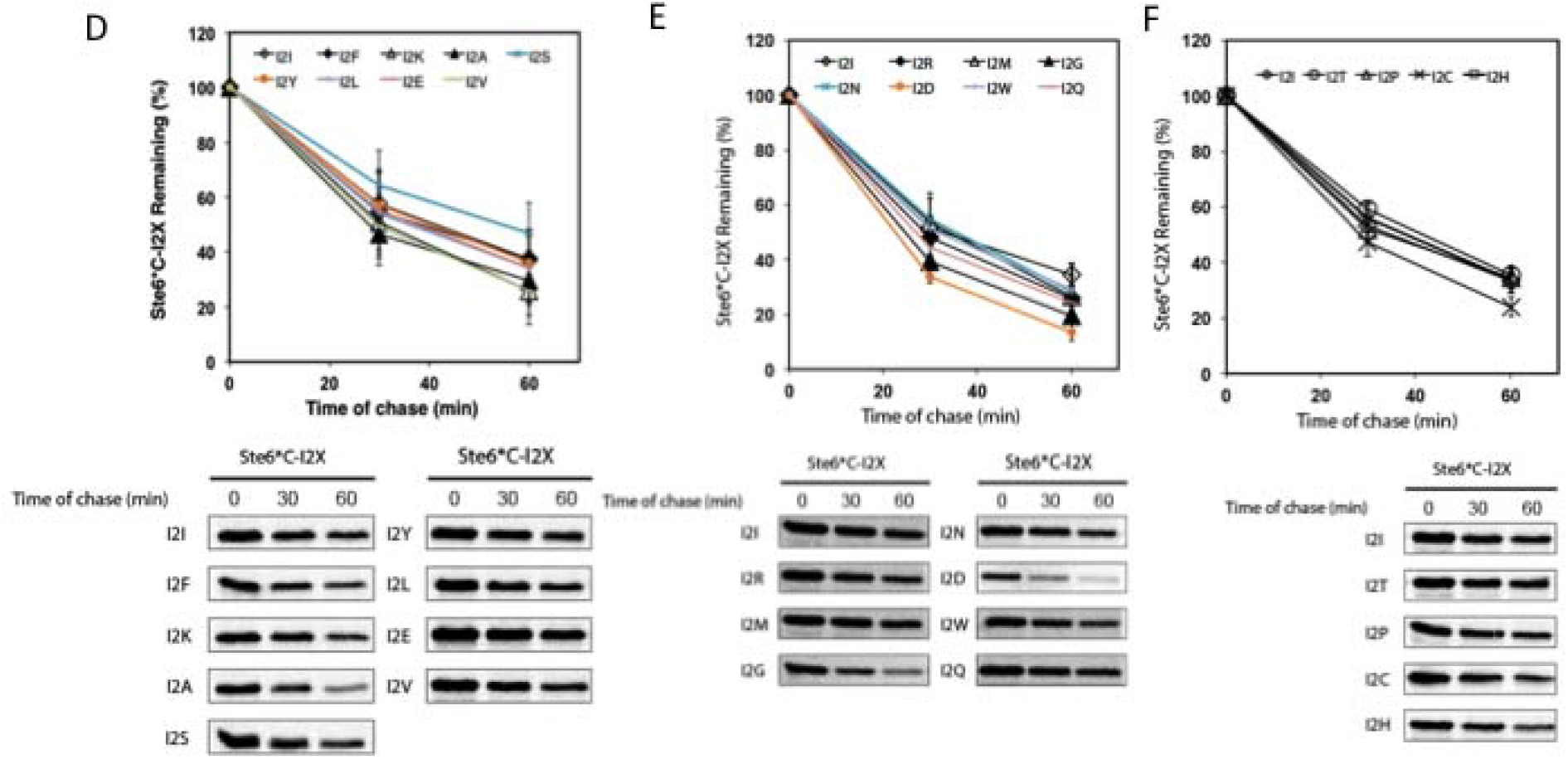
Efficient degradation of Ste6*C by the Ubr1p quality control pathway is highly ly dependent on P2 residue identity. (A), (B), and (C) Turnover rates of nineteen Ste6*C P2 P2 amino acid residue mutants (Ste6*C-I2X, where X = A[alanine], N[asparagine], S[serine],, C[cysteine], E[glutamic acid], P[proline], D[aspartic acid], T[threonine], M[methionine], H[histidine], W[tryptophan], F[phenylalanine], Y[tyrosine], V[valine], Q[glutamine],, G[glycine], K[lysine], L[leucine], R[arginine]) was compared to the turnover rate of the original al Ste6*C substrate (Ste6*C-I2I) in Δsan1 + UBR1 background cells by pulse chase analysis as as described in Fig 1A. (D), (E) and (F) Same as described in S2A, but in + SAN1Δubr1 cells. Error ror bars, mean +/- SD of three independent experiments (N = 3, biological replicates) unless ss otherwise noted. Due to an N = 2 for Ste6*C-I2 K expressed in + SAN1Δubr1 cells because of contamination (Fig S2D), lack of change vs Ste6*C-I2I could not be statistically confirmed; however, in both biological replicates of the experiment, the Ste6*C-I2 K substrate was degraded to a further extent than Ste6*C-I2I,indicating that the I2 K mutation does not inhibit degradation.

**Fig. S3.**
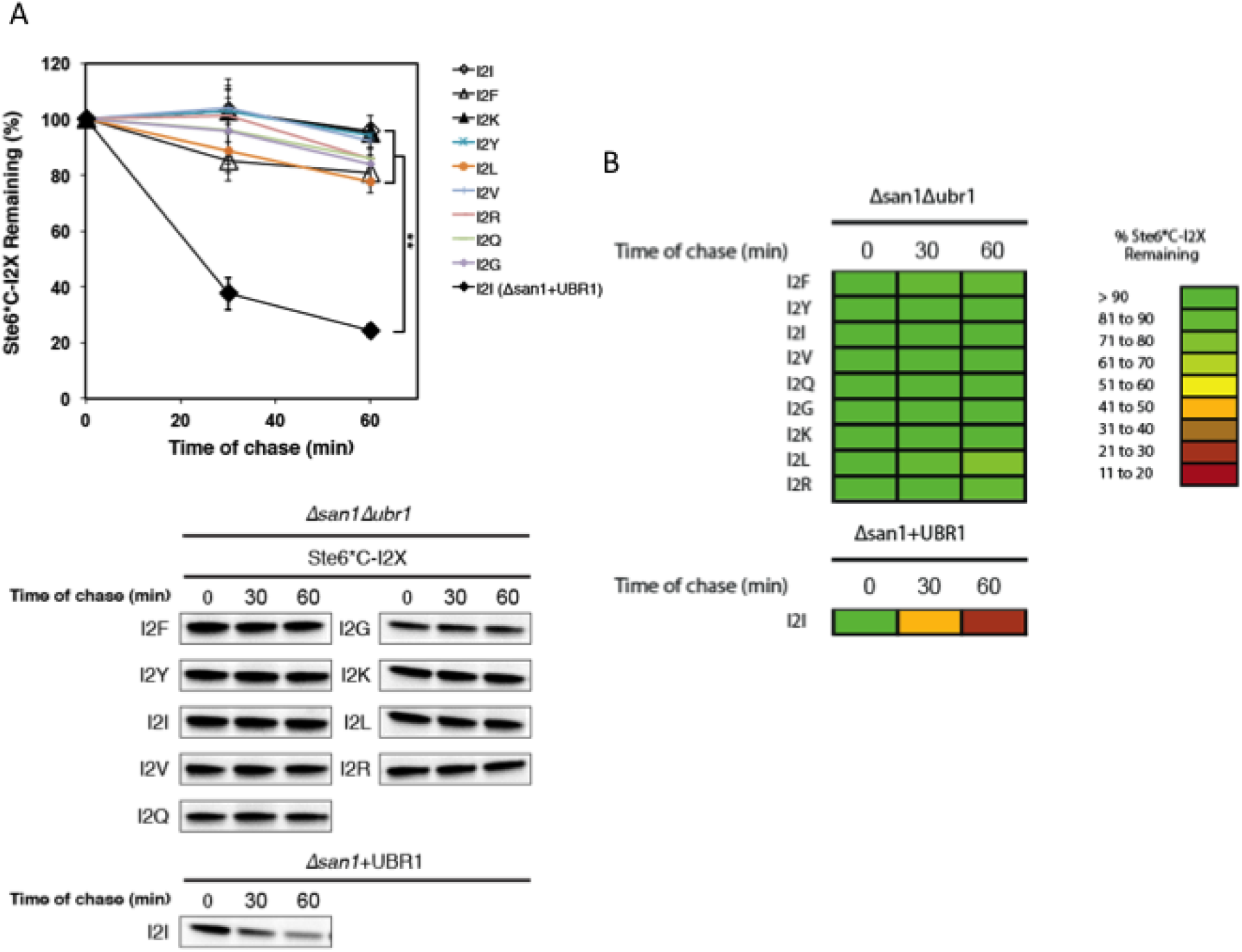
Ste6*C-I2X mutants bearing destabilizing P2 residues are stabilized in Δsan1Δubr1 cells. (A) Turnover rates of N-terminally destabilized Ste6*C-I2X mutants in Δsan1Δubr1 cells lls were assessed by pulse-chase analysis as described in Fig 1A. Degradation of Ste6*C-I2I in in Δsan1 + UBR1 cells was also examined as a control. Error bars, mean +/-SD of three independentnt experiments (N = 3, biological replicates). (B) Heat map reflecting turnover rates of Ste6*C-I2X X substrates in Δsan1Δubr1 as presented in S3A.

**Fig. S4.**
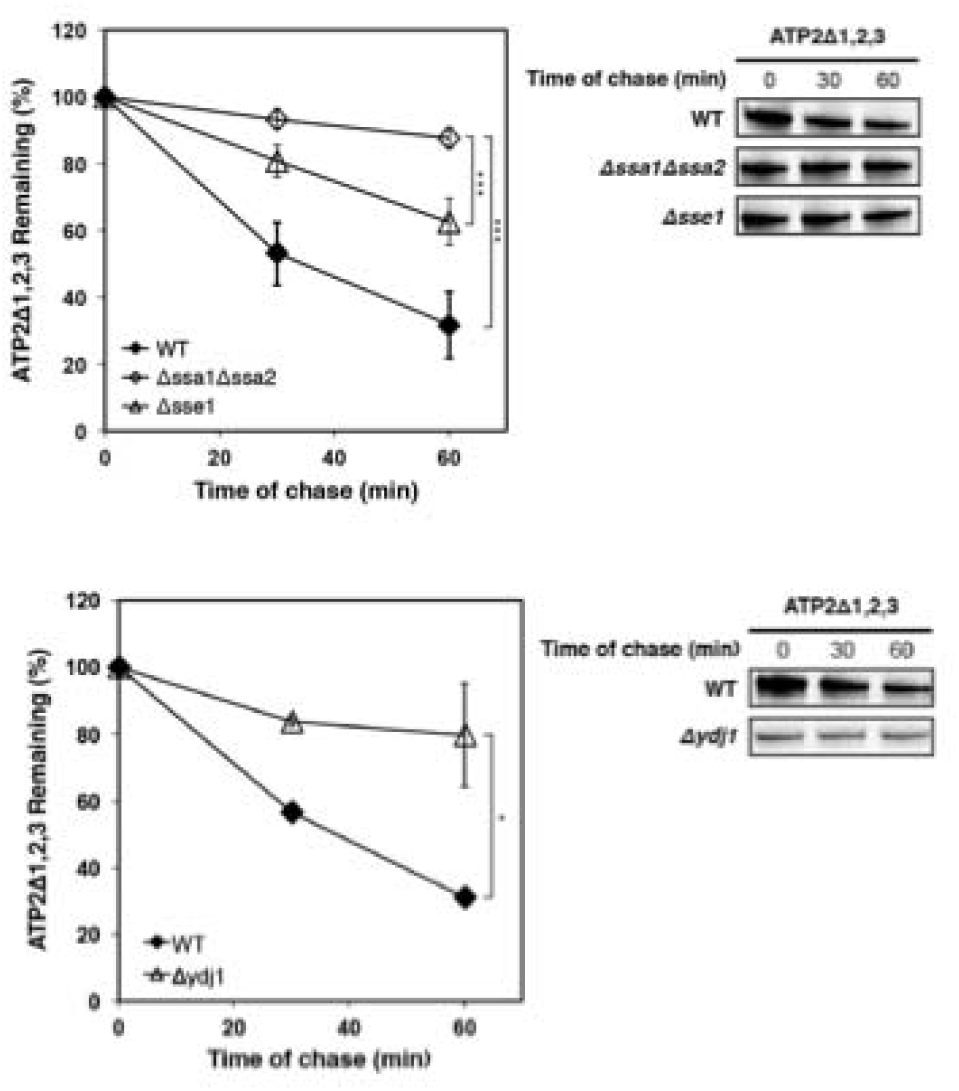
Degradation of ATP2Δ1,2,3 is dependent on chaperones involved in CytoQC C degradation. Turnover of ATP2Δ1,2,3 in WT, Δydj1, Δsse1Δsse2, and Δssa1 cells was analyzed ed by pulse-chase as described in in 5A. Student’s t-test: *, p < 0.05; **, p < 0.01; ***, p < 0.005; 5; Not Significant (NS), p > 0.5. Error bars, mean +/- SD of three independent experiments (N = 3,3, biological replicates).

**Fig. S5.**
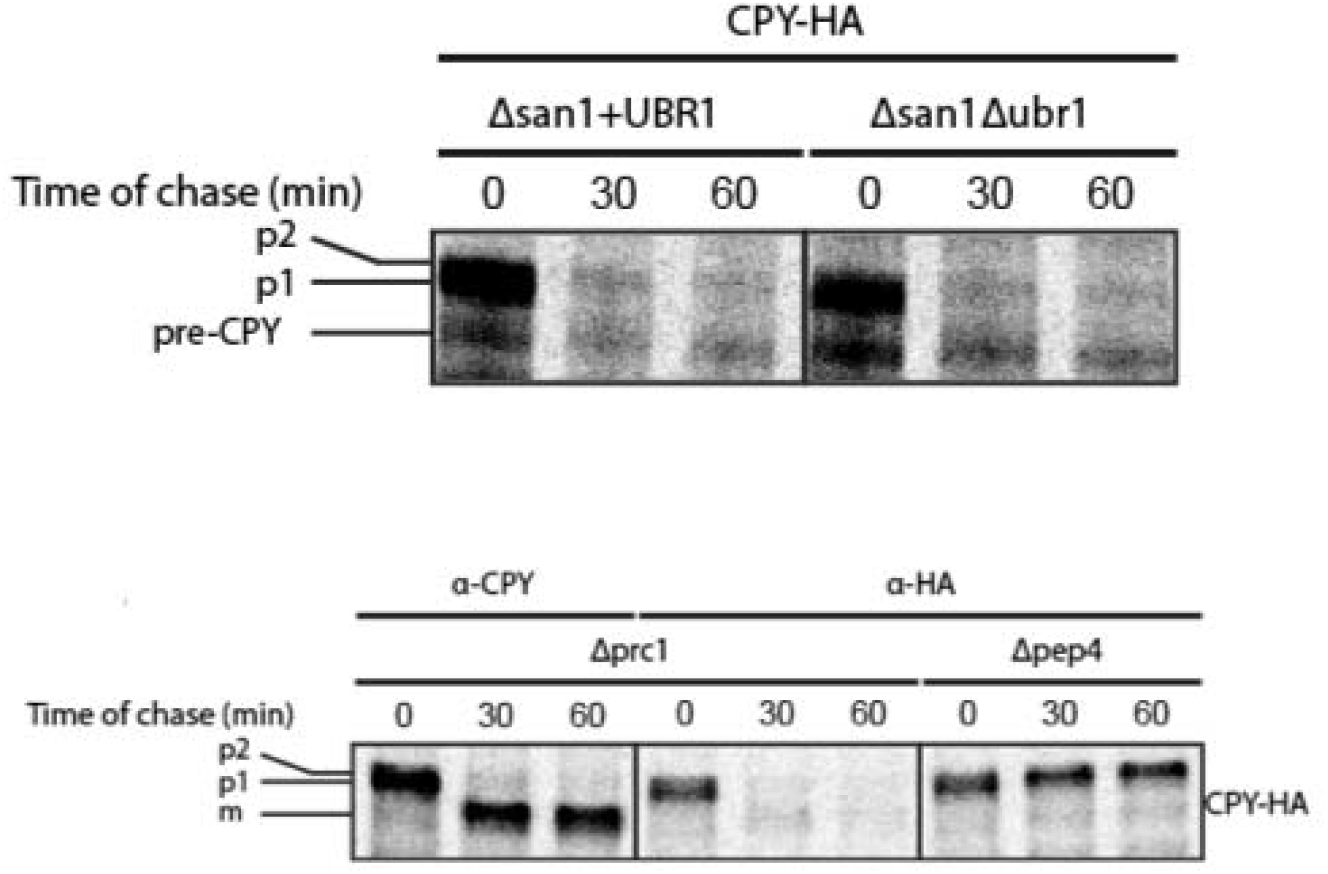
C-terminal HA-tag is removed from HA-tagged CPY in the vacuole. Pulse-chase analysis of CPY-HA was performed in Δprc1, Δpep4, Δsan1 + UBR1, and Δsan1Δubr1 cells as described in Fig 1A. CPY was probed with anti-CPY and anti-HA antibodies in Δprc1 cells, and anti-HA in Δpep4 cells.

